# Co-inhibition of topoisomerase 1 and BRD4-mediated pause release selectively kills pancreatic cancer *via* readthrough transcription

**DOI:** 10.1101/2023.02.10.527824

**Authors:** Donald P. Cameron, Jan Grosser, Swetlana Ladigan, Vladislav Kuzin, Evanthia Iliopoulou, Anika Wiegard, Hajar Benredjem, Sven T. Liffers, Smiths Lueong, Phyllis F. Cheung, Deepak Vangala, Michael Pohl, Richard Viebahn, Christian Teschendorf, Heiner Wolters, Selami Usta, Keyi Geng, Claudia Kutter, Marie Arsenian-Henriksson, Jens T. Siveke, Andrea Tannapfel, Wolff Schmiegel, Stephan A. Hahn, Laura Baranello

## Abstract

Pancreatic carcinoma is one of the most lethal cancers and the absence of efficient therapeutic strategies results in poor prognosis. Transcriptional dysregulation due to alterations in KRAS and MYC impacts initiation, development, and survival of this tumor type. Using patient-derived xenografts of pancreatic carcinoma driven by KRAS and MYC oncogenic transcription, we show that co-inhibition of Topoisomerase 1 (TOP1) and bromodomain containing protein 4 (BRD4) synergistically induce tumor regression through targeting promoter pause-release, a rate-limiting step in transcription elongation. By comparing the nascent transcriptome with the recruitment of elongation and termination factors along genes, we found that co-inhibition of TOP1 and BRD4, while globally impairing RNA production, disturbs recruitment of proteins involved in termination. Thus, RNA polymerases continue transcribing downstream of genes for hundreds of kilobases leading to readthrough transcription. This pervasive transcription also occurs during replication, perturbing replisome progression and leading to DNA damage. The synergistic effect of TOP1 and BRD4 inhibition is specific for cancer cells leaving normal cells unharmed, highlighting the sensitivity of the tumor to these transcriptional defects. This preclinical study provides a mechanistic understanding of the benefit of combining TOP1 and BRD4 inhibitors to treat pancreatic carcinomas addicted to oncogenic drivers of high transcription and replication.

**One Sentence Summary:** TOP1 and BRD4 inhibitors synergize to selectively kill pancreatic cancer *in vivo via* readthrough transcription without emergence of drug resistance

## INTRODUCTION

Dysregulated transcriptional programs cause cancer cells to become highly dependent on certain regulators of gene expression *(1–5)*. This dependency may provide new opportunities for novel therapeutic interventions. Pancreatic ductal adenocarcinoma (PDAC) is a highly lethal malignancy due to the lack of early diagnosis and limited response to treatments, with PDAC patients having a 5-year survival of 9% *(6, 7)*. Most PDACs harbor oncogenic KRAS mutations and elevated MYC signalling leading to dysregulation of global transcription and proliferation *(8–10)*, potentially sensitizing cancer cells to therapeutic targeting with transcriptional inhibitors.

Following transcription initiation, the RNA polymerase II (RNAPII) pauses due to factors that 1) affect the stability of the elongation complex and the efficiency of nucleotide incorporation *(11)*; and 2) provide a physical obstacle for the movement of RNAPII *(12, 13)*. The stages preceding and following pausing are associated with modifications of the RNAPII carboxyl-terminal domain (CTD). This CTD ‘‘code’’ defines mRNA splicing, elongation, histone methylation, and polyadenylation *via* an array of dynamic interactions *(14)*. The chromatin reader bromodomain containing protein 4 (BRD4) facilitates pause-release via two independent pathways: by recruiting the positive transcription elongation factor beta (PTEF-b) complex on the RNAPII *(15)* and by enhancing the enzymatic activity of topoisomerase 1 (TOP1) *via* phosphorylation of RNAPII CTD on Serine-2 (Ser-2) *(16, 17)*. TOP1 removes supercoiling in the DNA by transiently breaking one strand to allow controlled DNA rotation around the unbroken strand and resealing the break *(18)*. According to the ‘twin domain model’ *(19)*, as DNA twists through the active site of an elongating RNAPII, positive supercoils are generated ahead and negative supercoils trail behind the polymerase. Unless removed, this supercoiling will eventually halt the RNAPII. We have previously discovered a mechanism through which the RNAPII regulates TOP1 activity to favor transcription elongation. A positive feedback loop is established between RNAPII and TOP1 through their physical interaction. Upon BRD4-dependent phosphorylation *(17)*, the RNAPII-CTD directly stimulates TOP1 beyond its intrinsic activity to remove the supercoiling that would otherwise oppose pause-release *(16)*. BRD4-stimulated RNAPII-CTD phosphorylation is also reported to be involved in the recruitment of transcription termination factors (TTFs) to facilitate timely termination *(20)*. These TTFs, including cleavage and polyadenylation specificity factor (CPSF) and 64 kDa cleavage stimulation factor (CstF64), are loaded onto RNAPII in a BRD4-dependent manner at the 3’ end of genes. CPSF and CstF64 then promote the dephosphorylation of the elongation factor Spt5 by the Protein Phosphatase 1 (PP1) Nuclear Targeting Subunit (PNUTS) complex, thus slowing the RNAPII and enabling DNA disengagement *(21)*.

Although BRD4 belongs to the bromodomain and extra-terminal domain (BET) family—a group of proteins known to interact with acetylated histones—the stimulation of TOP1 activity by BRD4 *via* RNAPII-CTD phosphorylation is independent of nucleosome acetylation *(16)*. Thus, BRD4 acts through a bromodomain-dependent arm to drive PTEF-b on the pausing site, and through a bromodomain-independent arm to stimulate TOP1. The simultaneous targeting of BRD4 and TOP1 with a panel of BET inhibitors and TOP1 poisons respectively, synergistically inhibited cancer cell growth *in vitro (16)*. Thus, the combined inhibition of both arms could be used therapeutically to target the transcriptional addiction of PDACs dependent on KRAS and MYC dysregulated gene expression programs *(2)*.

Here we tested the efficacy of this combinatorial strategy *in vivo* using a collection of patient-derived xenograft (PDX) mouse models for PDAC. We also included a pancreatic neuroendocrine carcinoma (panNEC) harboring a KRAS mutation and elevated MYCN, the MYC isoform expressed in neurons. PanNECs represent a poorly differentiated and highly malignant subgroup of neoplasms of the neuroendocrine pancreas. Although panNECs are rare malignancies *(22, 23)*, their incidence has increased steadily, especially as metastatic disease *(24, 25)*. The TOP1 poison Irinotecan is already used to treat PDACs and panNECs as part of the FOLFIRINOX and FOLFIRI drug cocktails, respectively *(26, 27)*. However, while this treatment provided median overall survival of 11 months, it was associated with significant side effects reducing patient quality of life. Since synergistic drug combinations enable reduced dosage while retaining efficacy, combining TOP1 and BRD4 inhibitors might represent a promising strategy to pharmacologically uncouple the TOP1-RNAPII regulation of transcription elongation, while reducing toxic nonspecific DNA damage associated with classical TOP1 inhibitors *(28, 29)*. We show that combining the TOP1 inhibitor Irinotecan with the BRD4 inhibitor JQ1, synergistically kills the tumor cells in the PDX model of both tumor types. Mechanistically, we found that the inhibition of both bromodomain-dependent and independent pathways profoundly impairs promoter-proximal pausing, transcriptional elongation, and the downstream mechanisms of termination, leading to readthrough transcription. Thus, transcribing RNAPIIs remain engaged with DNA for hundreds of kilobases (kb) into largely gene-free late replication regions, causing replication stress and inducing DNA damage and cellular stress signaling. Our results demonstrate that the synergistic drug combination selectively affects tumor viability based on their transcriptional addiction, leaving normal cells largely unaffected.

## RESULTS

### Combined inhibition of BRD4 and TOP1 is synergistic in killing xenografted tumors of pancreatic cancer

We previously demonstrated that targeting two independent arms of promoter-proximal pausing regulation through inhibition of TOP1 and BRD4 (Fig. 1A) synergistically inhibited tumor growth *in vitro (16)*. To determine whether this combination could effectively arrest cancer progression *in vivo*, mice with implanted pancreatic cancer were treated with either the clinically approved TOP1 poison Irinotecan (15 mg/kg, three times weekly, every second week), the BRD4 inhibitor JQ1 (50 mg/kg, daily) or both drugs in combination. Notably, while the JQ1 dose is in accordance with other studies, the Irinotecan dose used in our study is considerably lower than common administration regimes of 40-300 mg/kg daily *(30, 31)* to limit the nonspecific cytotoxic effects associated with TOP1 drugs.

**Fig. 1.**
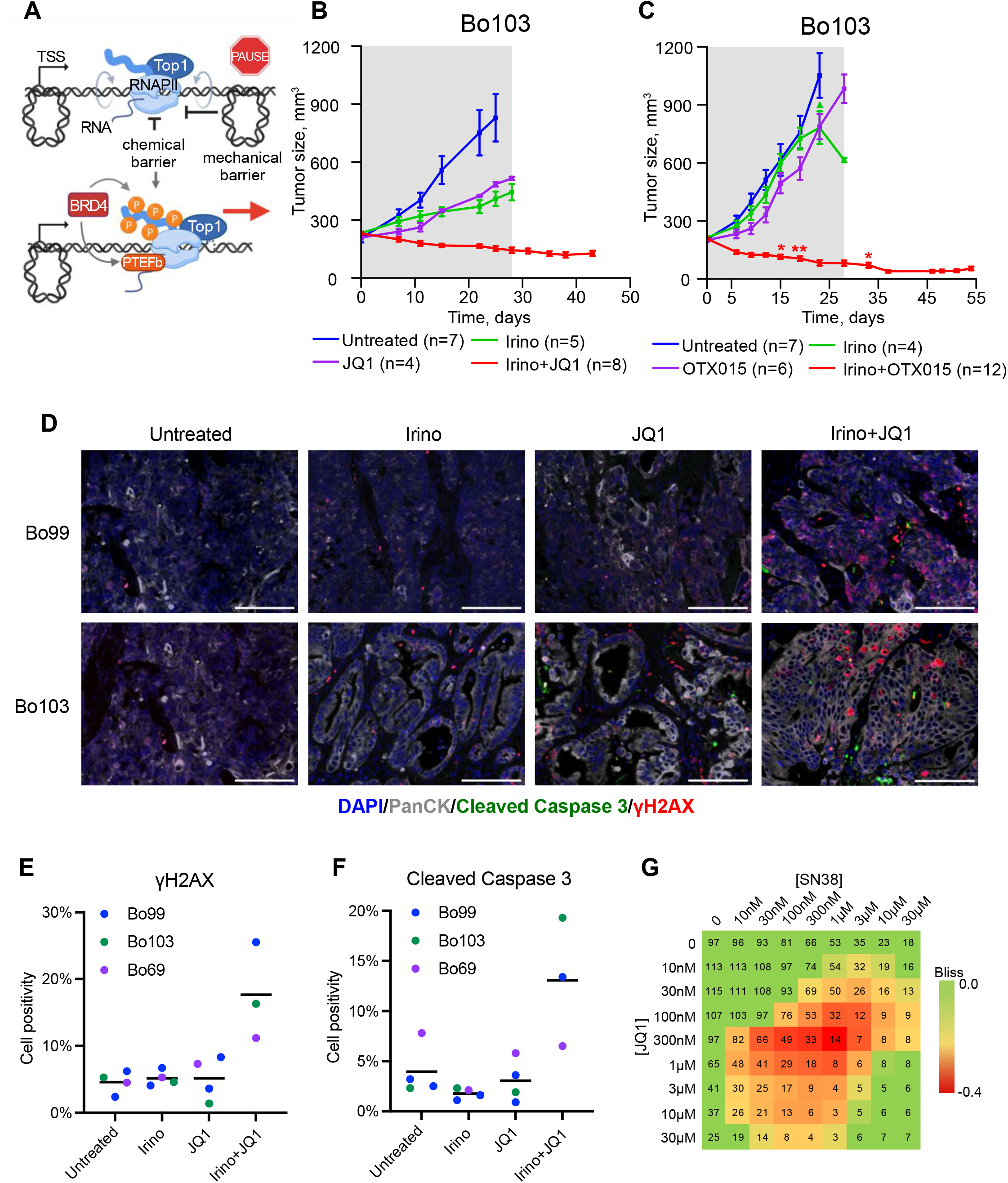
Combined drug treatment of TOP1 and BRD4 inhibitors shows synergy by killing pancreatic tumor cells both *in vivo* and *in vitro*. **(A)** Scheme adapted from Baranello et al. *(16)* showing that BRD4 and TOP1 activity are required to overcome promoter-proximal pausing and enable efficient transcription elongation. **(B)** Primary responses observed within the 28 days treatment interval (shaded area) for the PDAC PDX models Bo103 treated with Irinotecan (Irino) (15 mg/kg, three times weekly, every second week) and JQ1 (50 mg/kg, daily) by intraperitoneal (i.p.) injection, alone or in combination in comparison to untreated controls. Growth curves are derived from mean values ± SEM (error bars). **(C)** Same as (B), but with the BRD4 inhibitor OTX015. Each asterisk represents a mouse that was taken out of the treatment cohort at the indicated time point because of health issues of the animal. The triangle indicates a mouse taken out of the experiment because one of the two tumors reached the maximum size criteria. Two mice with altogether 4 tumors were kept for follow-up beyond end of treatment on day 28 to assess tumor recurrency. **(D)** Representative immunohistochemistry images of Bo99 and Bo103 PDX tumor sections treated for 5 days with Irinotecan and/or JQ1 stained for DNA (DAPI, blue), PanCK (grey), cleaved caspase 3 (green) and γH2AX (red). Scale bar represents 100 μm. **(E-F)** Quantitation of nuclear γH2AX (E) and cellular cleaved caspase 3 (F) positivity from samples in (D) and fig. S1D. **(G)** Checkerboard assay of cultured Bo103 cells treated with increasing concentrations of SN38 and JQ1 in combination as indicated. The percentage of confluency after treatment is denoted by the numbers in the squares. Synergy was determined using the delta Bliss model of additivity with lower, more negative values showing stronger synergy, (visualized by red/green color coding). Representative checkerboard of n=3.

Four PDX models with activated oncogenic KRAS mutations and elevated MYC isoforms (table S1), known to dysregulate global transcription *(8, 9)*, were chosen. Each monotherapy was able to induce a certain degree of tumor growth reduction without inducing tumor regression (Fig. 1B, and fig. S1, A to C). However, the combination therapy induced detectable tumor regression (mean maximal tumor volume reduction of on average 32-49% relative to the mean tumor volume at therapy start) after 28 days in the three PDAC PDX models, including one PDAC model (Bo103) belonging to the classical subtype and two (Bo69 and Bo85) belonging to the quasi*-*mesenchymal subtype *(32)*, the latter subtype being frequently linked to therapeutic resistance. Of note, the panNEC PDX model (Bo99) showed a near 100% tumor volume reduction. The synergism was also prominent when Irinotecan was combined with OTX015, a JQ1 analog and clinical stage bromodomain inhibitor, indicating this drug targeting strategy could prove promising in cancer patients (Fig. 1C). Immunohistochemistry staining of Bo69, Bo99 and Bo103 tumor sections demonstrated an increase in DNA damage response marker γH2AX and apoptotic marker cleaved caspase 3 after Irinotecan+JQ1 treatment (Fig. 1, D to F, and fig. S1D). This effect was most pronounced in the Bo99 and Bo103 tumors, as the Bo69 section had high cleaved caspase 3 background signal. Given the Irinotecan treatment alone showed no increase in γH2AX relative to the untreated samples (Fig. 1E), the DNA damage signaling must be specific to the combination treatment, as opposed to the genotoxic properties of Irinotecan. These *in vivo* experiments strongly indicate that the Irinotecan+JQ1 treatment synergistically induces targeted DNA damage, apoptosis, and tumor shrinkage.

Of the tumors tested, Bo103 was the most adaptable for cell culture. We confirmed that the *in vitro* co-inhibition of BRD4 and TOP1 with JQ1 and SN38, the active metabolite of Irinotecan required for cell culture *(33, 34)*, was synergistic in killing cells as determined by the Bliss independence model of additivity *(35)*, while no synergy was observed in the normal immortalized pancreatic cell line hTERT-HPNE (Fig. 1G and fig. S1F). Approximately 90% growth inhibition was detected after 48 hr of combined treatment with 500 nM SN38 and 1 μM JQ1, whereas individual administration had limited growth inhibitory effects (Fig. 1G and fig. S1E). Therefore, the Bo103 cells and these drug concentrations were subsequently used to characterize the tumor-specific mechanisms underlying the synergy.

### SN38 and JQ1 combination treatment synergistically inhibits transcription

To assess the effect of SN38 and JQ1 on TOP1 activity and BRD4 localization respectively, we performed TOP1 Covalent Adduct Detection-sequencing, or TOP1 CAD-Seq *(36, 37)* and BRD4 chromatin immunoprecipitation sequencing with spike-in control, or ChIP-Rx-Seq *(38)*. TOP1 CAD-Seq enables quantification of catalytically active TOP1 on the DNA, known as the TOP1 cleavage complex (TOP1cc). This complex is extremely transient in cells, unless stabilized with TOP1 poisons and inhibitors of proteasomal degradation *(39)*. Bo103 cells were treated with vehicle, SN38 or SN38+JQ1 for 1 hr. For the final 30 min, the proteasome inhibitor MG132 was added to all conditions to inhibit degradation of the Top1ccs *(40)*. SN38 treatment resulted in increased detection of TOP1ccs downstream of the TSS where the phosphorylated CTD of RNAPII stimulates TOP1 activity relative to the untreated condition *(16)* (Fig. 2A). Notably, we found proportionally fewer TOP1ccs towards the transcription end site as compared to the TOP1cc profile measured across the genes upon shorter (4 min) Irinotecan analog Camptothecin treatment *(16)*. This likely happens because TOP1 inhibition blocks RNAPII elongation *(41)*, leading to fewer TOP1ccs engaged across the gene body. Upon treatment with SN38+JQ1, the portion of TOP1ccs covalently engaged with the DNA downstream of the TSS were reduced, as compared to SN38 alone likely reflecting the reduced stimulation of TOP1 activity due to a decrease in BRD4-dependent CTD phosphorylation *(17)* (Fig. 2A). Indeed, the BRD4 ChIP-Rx-Seq experiment confirmed that while SN38 increased BRD4 at the TSS likely due to trapped TOP1 blocking BRD4 release from the promoter, JQ1 alone or in combination with SN38 reduced BRD4 occupancy (Fig. 2B). Because BRD4 and TOP1 are key factors in the regulation of promoter-enhancer activity *(42, 43)*, their loss is expected to severely affect enhancer function, as measured by the active enhancer marker H3 lysine 27 acetylation (H3K27Ac). Indeed, H3K27Ac ChIP-Rx-Seq revealed a profound decrease in lysine 27 acetylation only upon combination treatment (Fig. 2C and fig. S2A). This decrease was evident both at enhancers and at promoters, indicating that SN38+JQ1 incubation globally impairs transcription.

**Fig. 2.**
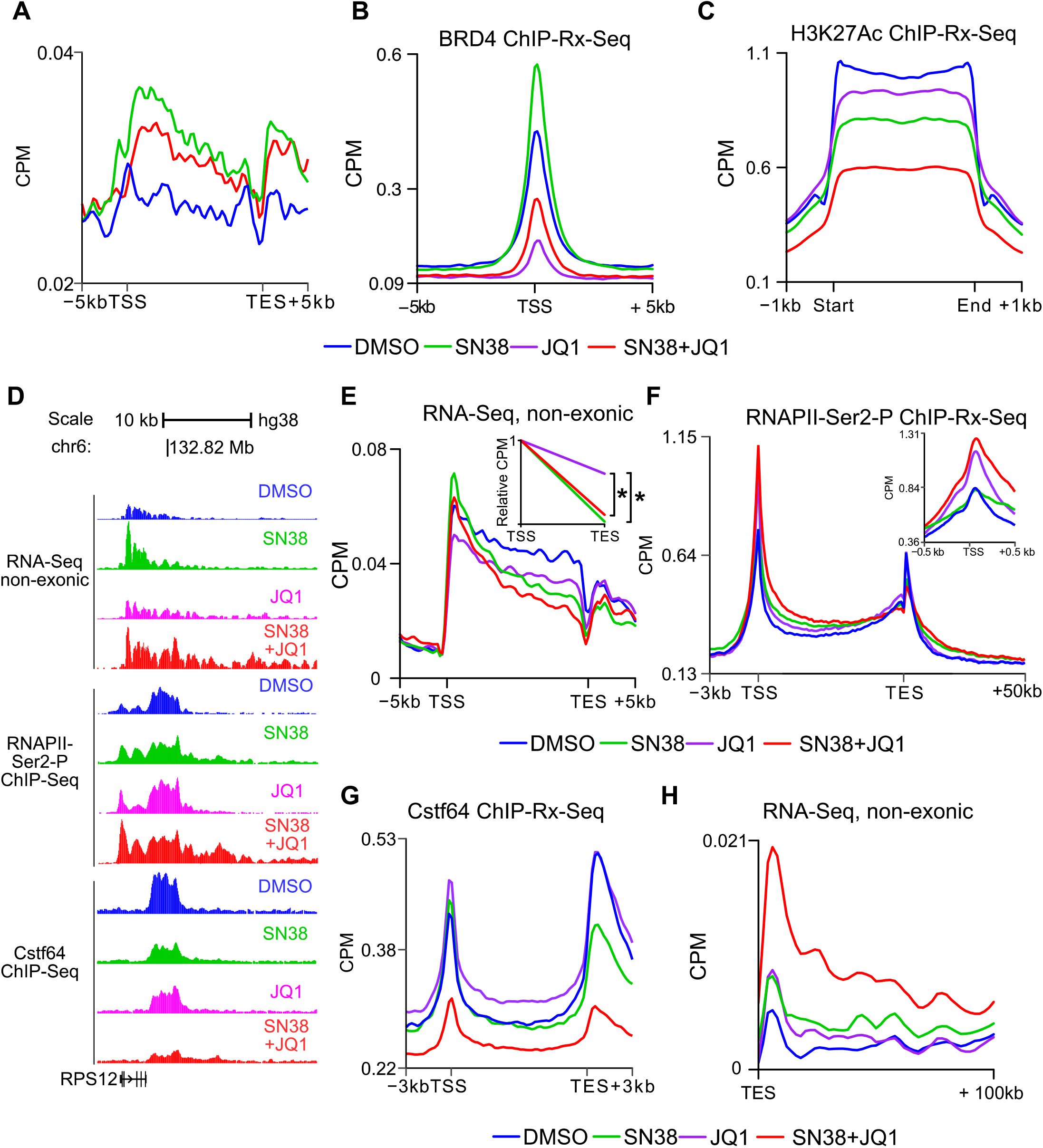
Transcription is synergistically inhibited by the combination treatment SN38+JQ1. **(A)** TOP1 CAD-Seq profile at the 2,500 most expressed genes between transcription start site (TSS) and transcription end site (TES) in Bo103 cells treated with DMSO, SN38 or SN38+JQ1 for 1 hr. Data represented as count per million reads (CPM). Average of biological duplicates. MG132 only treatment showed similar profile to vehicle, excluded for clarity. **(B)** BRD4 occupancy at the TSS of the 10,000 most expressed genes (+/-5 kb) in Bo103 cells after 4 hr. Data are spike-in normalized. Average of biological duplicates. **(C)** H3K27ac occupancy at enhancers (+/-1 kb) predicted from H3K27ac peak calling*(42, 102)* in Bo103 cells treated with DMSO, SN38, JQ1, or SN38+JQ1 for 4 hr. Data are spike-in normalized. Average of biological duplicates. **(D)** Example Genome Browser tracts of non-exonic RNA-Seq, Cstf64 and RNAPII-Ser2-P ChIP-Rx-Seq reads along the gene body and downstream of the gene RPS12 for all treatment conditions after 4 hr. **(E)** Non-exonic RNA-Seq reads from Bo103 cells plotted between TSS and TES of the 10% longest protein-coding genes after 4 hr of treatment. Inset shows the gradient of the linear regression between TSS and TES (NERD index; DMSO and JQ1 are indistinguishable). Average of biological triplicates. *: p<0.05, Student’s t-test. **(F)** RNAPII-Ser2-P occupancy at the 10,000 most expressed genes in Bo103 cells. Data are spike-in normalized. Inset shows the distribution of RNAPII-Ser2-P around TSS after 4 hr. CPM range differs between main figure and inset due to differential binning based on gene proportion and absolute bp number, respectively. Average of biological duplicates. **(G)** Cstf64 occupancy at the 10,000 most expressed genes in Bo103 cells after 4 hr. Spike-in normalized. Average of biological duplicates. **(H)** Non-exonic RNA-Seq reads from Bo103 cells plotted in the 100 kb downstream of the TES of protein-coding genes after 4hr. Average of biological triplicates.

Our model (Fig. 1A) predicts that inhibiting BRD4 recruitment on chromatin and TOP1 stimulation will prevent RNAPII pause-release, thus impairing RNAPII progression. If true, this would result in accumulation of short RNA species around the beginning of genes and reduced amount of RNA in the end of the genes. To test this hypothesis, we did RNA-seq performing the step of retro-transcription with random primers and plotted only non-exonic reads on the metagene (Fig. 2D and fig. S2B). These reads correspond to regions that are typically excised and degraded during co-transcriptional splicing *(44)*, allowing for determination of nascent transcription. Upon 4 hr SN38 treatment, the profile of nascent transcripts paralleled the TOP1 CAD-Seq (Fig. 2A) showing a buildup of transcripts at the beginning of genes with a depletion of RNAs towards the ends of genes (Fig. 2D, fig. S2B, and table S2), most pronounced in long genes (Fig. 2E). The extent of this change can be quantified using the gradient of the linear regression of the read distribution, which we term the Non-Exonic Read Distribution index, or NERD index (Fig. 2E, inset, and fig. S2B, inset). Although the NERD index after JQ1 treatment was similar to DMSO, we found an overall reduction in nascent transcripts across the entire gene unit, suggesting defects in RNAPII initiation or elongation. This was possibly due to inhibition of BRD4 bromodomain interaction with acetylated histones at promoters of genes *(45)*. The SN38+JQ1 treatment displayed both a decreased NERD index and reduced number of nascent transcripts, indicating that RNAPII transcription was targeted independently through both pathways (Fig. 1A). These trends were reproducible by SLAM-seq *(46)*, where nascent transcripts are directly labelled, indicating that the results do not simply reflect altered splicing (fig. S2C). The SN38+JQ1 treatment also exhibits more differentially expressed genes relative to individually treated samples (fig. S2D), with “Transcription by RNA polymerase II” and “Regulation of gene expression” among the top downregulated gene ontologies, indicating that the cells might undergo reprogramming to reduce overall transcription (fig. S2E). Thus, the SN38+JQ1 seemed to affect transcriptional pause release and subsequent elongation, as predicted, resulting in stronger cellular response than each individual drug.

### Transcription termination is synergistically inhibited by SN38+JQ1 treatment

BRD4 controls RNAPII transcription by modulation of RNAPII CTD phosphorylation and TTF recruitment *(47, 48)*. Given the loss of promoter-bound BRD4 observed with JQ1 alone or in combination with SN38 (Fig. 2B), that would lead to an altered pattern of RNAPII Ser-2 phosphorylation (Ser2-P) and reduced TTF engagement *(20)*. ChIP-Rx-Seq of RNAPII Ser2-P upon JQ1 treatment showed an increase in Ser2-P levels at promoter-proximal regions (Fig. 2F), similar to previous findings upon BET protein degradation *(20)*. Treatment with SN38+JQ1 exaggerated this effect (Fig. 2F). This might represent a futile attempt of the system to compensate for the absence of pause-release regulation, by releasing stored PTEF-b *(49)* and delivering it to the RNAPII *(50)*. Surprisingly, the RNAPII Ser-2P signal also extended far downstream the transcription end site (TES) after SN38+JQ1 when compared to the untreated control, suggesting potential defects in transcription termination (Fig. 2, D and F). In accordance, ChIP-Rx-Seq showed that the binding of the 3’ RNA processing factor Cstf64 at both the TSS and TES was markedly reduced after SN38+JQ1 treatment (Fig. 2, D and G).

Aberrant recruitment of 3’ RNA processing factors have been associated with transcription readthrough of RNAPII downstream the 3’ end of genes *(20)*. Analysis of the non-exonic reads from the RNA-Seq revealed that SN38+JQ1 led to an increase in RNA signal downstream of the 3’ end extending even beyond 100 kb (Fig. 2H and fig. S2B). These Downstream of Gene (DoG) transcripts were clearly elevated after the combination treatment relative to all other conditions (fig. S2F). Together, these data suggest that SN38+JQ1 act synergistically to impair transcription, while paradoxically leading to readthrough transcription far downstream of the TES due to loss of recruitment of termination factors.

### The genes exhibiting readthrough transcription are highly expressed and heavily paused

Although Cstf64 binding was reduced in all genes after SN38+JQ1, DoG transcription was only detected in a subset of genes. To further characterize why DoG genes are vulnerable to readthrough transcription, we generated a set of non-DoG genes of similar length and expression for comparison (Fig. 3A). Analysis of RNA-Seq downstream of the TES demonstrates that even in untreated conditions DoG genes exhibit higher levels of readthrough transcription compared to the non-DoG genes (Fig. 3B) suggesting that DoG are prone to exhibit readthrough transcription even at a basal state. This effect is accentuated after SN38+JQ1 exposure (Fig. 3B). Because of their dependency on TOP1 activity and BRD4 regulation, we surmised that genes undergoing readthrough transcription must be highly expressed and highly paused. Indeed, we observed that most genes exhibiting readthrough transcription were characterized on average by higher levels of expression in comparison to all expressed genes (fig. S3A). Also, while comparison of these two gene sets demonstrated that both have similar levels of Cstf64 loss after SN38+JQ1 treatment (Fig. 3C), DoG genes have a higher pausing index than non-DoG genes based on published RNAPII ChIP-Seq data *(51)* (Fig. 3D). Moreover, the DoG genes exhibited higher levels of RNAPII Ser-2P than non-DoG genes, suggesting stronger dependence on RNAPII CTD modifications (fig. S3B).

**Fig. 3.**
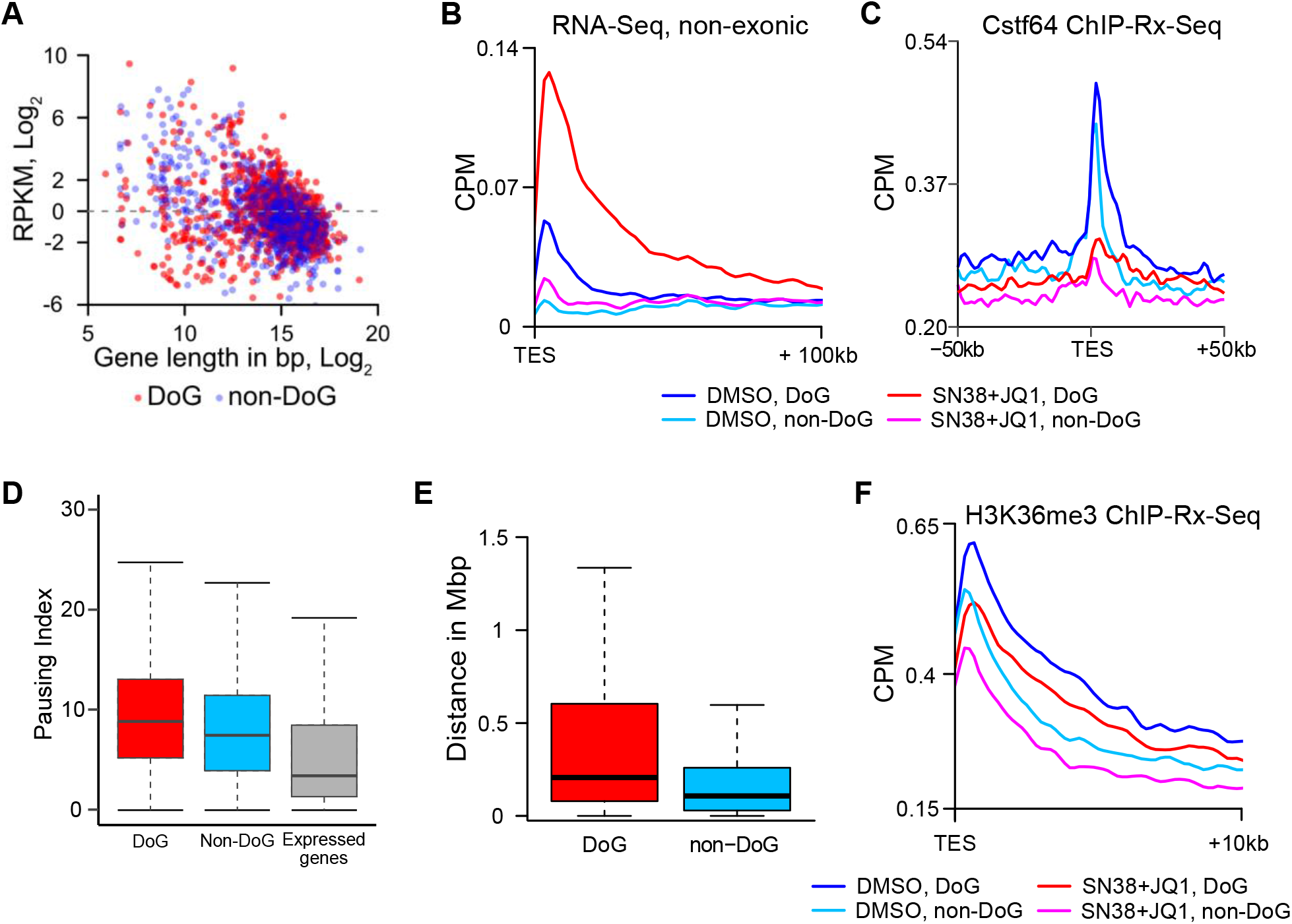
Readthrough transcription is associated with high gene expression and RNAPII pausing. **(A)** Comparison of expression level as reads per kilobase per million (RPKM) and gene length of detected genes producing DoG transcript after SN38+JQ1 treatment and the generated list of random non-DoGs. **(B)** Non-exonic RNA-Seq reads plotted for the region 100 kb downstream of the TES of DoG and non-DoG genes in untreated (DMSO) or SN38+JQ1 treated (4 hr) Bo103 cells. Average of biological triplicates. **(C)** Cstf64 occupancy around TES of DoG and non-DoG genes in Bo103 cells after 4 hr. Data are spike-in normalized. Average of biological duplicates. **(D)** Boxplot showing RNAPII pausing index of DoG, non-DoG and all expressed genes. Whiskers indicate lowest and highest values no further than 1.5 x interquartile range. **(E)** Boxplot showing the distance from the TES of high stringency DoGs and non-DoGs to the next expressed gene. Whiskers indicate lowest and highest values no further than 1.5 x interquartile range; outliers are excluded. **(F)** H3K36me3 occupancy 10 kb downstream of the TES of DoG and non-DoG genes in Bo103 cells after 4 hr treatment with DMSO or SN38+JQ1. Data are spike-in normalized. Average of biological duplicates.

If susceptibility to readthrough transcription originates from dysregulation of RNAPII modifications at the pausing site where TOP1 and BRD4 are functional, then readthrough would not be detected until the RNAPII had transcribed the length of the gene. Indeed, a qPCR time course experiment demonstrated that short genes like SERP1 (4.5 kb) and SEC61B (8.3 kb) showed readthrough transcription after 1 hr, while the longer gene BPNT2 (36 kb) did not exhibit readthrough transcription until 2 hr post-treatment with SN38+JQ1, despite the primers being equidistant from the respective TESs (fig. S3C). In addition, considering the average RNAPII transcription rate in the presence of TOP1 poison Camptothecin is 1 kb/min *(52)*, genes longer than approximately 240 kb should not exhibit readthrough transcription after 4 hr treatment. Indeed, DoG genes are typically shorter than 150kb in length (fig. S3D). These data support the concept that polymerases are primed to undergo readthrough transcription at the pausing site as opposed to the TES.

We reasoned that the readthrough transcription that extends hundreds of kb downstream the TES would likely disturb other chromatin-related processes. To this end, we curated a “high-stringency” list of DoG genes characterized by elevated transcription 30-45 kb downstream of the TES after SN38+JQ1 relative to untreated control. This subset was even more specific for SN38+JQ1, as the individual treatments had fewer DoG genes that matched these criteria (fig. S3E), and readthrough transcription could be detected up to 200 kb downstream of the TES (fig. S3F). To understand how this readthrough transcription may affect other intergenic or intragenic regions, we calculated the distance of the closest expressed gene downstream of the high-stringency DoG genes. Interestingly, readthrough transcription from these high-stringency genes was more likely to extend into gene-free regions relative to a non-DoG gene set of similar expression and length distribution as evident by the on average higher distance to the next expressed gene located downstream (Fig. 3E).

### Readthrough transcription affects repressive chromatin markers

If DoG transcripts are more likely to extend into gene free regions, their chromatin must be silenced to safeguard transcription fidelity. Lysine 36 of Histone 3 is typically tri-methylated (H3K36me3) by SET Domain Containing 2, Histone Lysine Methyltransferase (Setd2) following RNAPII passage to prevent spurious transcription *(53–55)*. In addition, loss of Setd2 is associated with increased readthrough transcription, suggesting H3K36me3 plays a role in blocking DoG transcripts *(56)*. As SN38+JQ1 treatment was found to reduce Setd2 expression (table S2) and Setd2 is the sole methyltransferase responsible for H3K36me3, we investigated this chromatin modification by ChIP-Rx-Seq. The aim of this experiment was to understand: 1) why genes show differential susceptibility to DoG transcription despite similar loss in Cstf64 binding; and 2) how induction of readthrough transcription by SN38+JQ1 treatment modulates the local chromatin environment. In untreated conditions, H3K36me3 levels were higher in the 10 kb region downstream of the 3’ ends of DoG *vs*. non-DoG genes (Fig. 3F, compare dark blue and light blue curves). Treatment with SN38+JQ1 overall reduced the amount of the H3K36me3 marker. Thus, the combined inhibition of TOP1 and BRD4 promotes changes in the chromatin structure, downstream of genes, towards a more de-repressed state.

### SN38+JQ1-driven readthrough transcription impacts late replication causing potential replication-transcription collisions

If the transcriptionally engaged RNAPIIs can continue transcribing into intergenic regions, thus remodeling the chromatin in those regions, they might potentially affect the translocation of other DNA revolving machineries, such as replisomes. To test this hypothesis, we first verified whether readthrough transcription is detectable during S-phase upon SN38+JQ1 treatment, and then whether it affects replication fork progression. Cells were synchronized in S-phase with the DNA synthesis inhibitor hydroxyurea (HU) *(57)*, released into fresh medium and treated with SN38 or JQ1 alone or in combination for 4 hr (fig. S4A). While in untreated conditions the RNA signal at the analyzed DoG regions was negligible, it increased remarkably upon drug treatment reaching its maximum with SN38+JQ1 (Fig. 4A, compare lanes 5 *vs*. 8). Importantly, we detected comparable DoG transcripts between S-phase-synchronized and asynchronous cells (Fig. 4A, compare lanes 4 *vs*. 8), indicating that SN38+JQ1-driven readthrough transcription persists during S-phase.

**Fig. 4.**
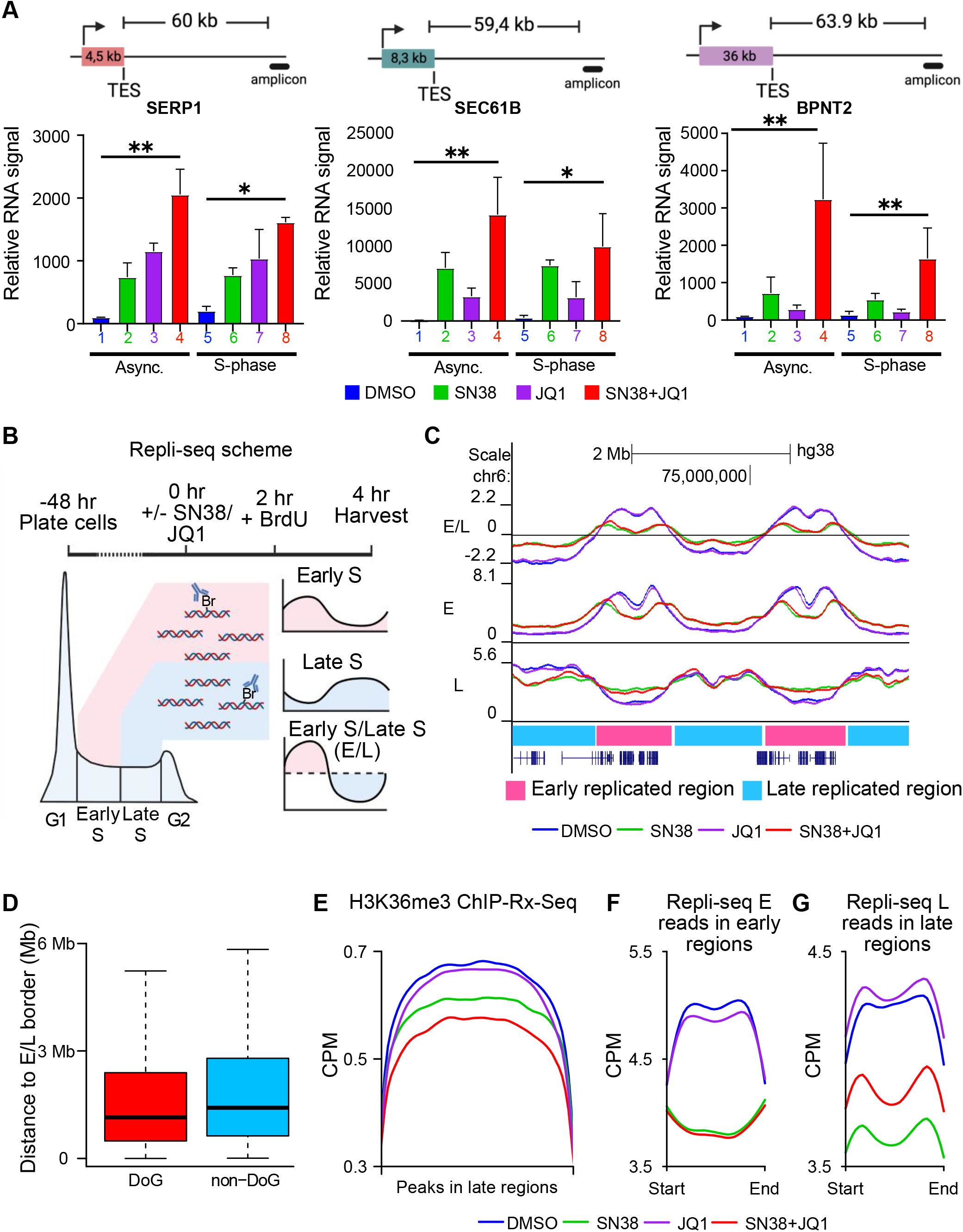
Readthrough transcription induced by SN38+JQ1 enhances firing of dormant origins by chromatin decompaction. **(A)** Readthrough transcription persists in S-phase upon treatment with SN38+JQ1, as detected at selected regions downstream of the DoG genes SERP1, SEC61B and BPNT2 after 4 hr (n=3, relative to DMSO control, error bars represent standard deviation). *: p<0.05; **: p<0.01, Student’s t-test. **(B)** Scheme of Repli-seq. The approach allows for determination of early and late replicated regions in the genome. **(C)** Example Genome Browser tracks of replication timing (E/L), early (E) and late (L) replication upon Bo103 cell treatment with DMSO, SN38, JQ1, or SN38+JQ1. Early (pink) and late (light blue) replicated regions are denoted by colored boxes underneath the tracks. Average of biological duplicates. **(D)** Boxplot showing the distance downstream from the TES of DoGs and non-DoGs in early replicated regions to the next early-to-late border (p-value < 0.0005). Whiskers indicate lowest and highest values no further than 1.5 x interquartile range; outliers are excluded. **(E)** H3K36me3 peaks in late replicated regions in Bo103 cells treated with DMSO, SN38, JQ1, or SN38+JQ1 for 4 hr. Data are spike-in normalized. Average of biological duplicates. **(F)** Repli-Seq E read coverage at early replicated regions. Average of biological duplicates. **(G)** Repli-Seq L read coverage at late replicated regions. Average of biological duplicates.

Next, we applied Repli-seq *(58, 59)* to observe changes in replication timing (RT) after 4 hr treatment with SN38 or JQ1, alone or in combination. Briefly, replicating cellular DNA was labelled by BrdU pulsing, cells were then harvested and sorted into early (E) and late (L) S-phase populations. BrdU-labelled DNA was then immunoprecipitated and sequenced to determine the replication timing (RT) of each genetic region (Fig. 4B). The RT (i.e. the regions referred to as early or late replicating based on the E/L profile) did not broadly change between treatment conditions, although the reduced signal in both SN38 and SN38+JQ1 conditions indicated that TOP1 inhibition greatly affects the overall replication rates (Fig. 4C), as expected, given the requirement of TOP1 for proper replication fork progression *(60)*.

We predicted that the readthrough RNAPIIs induced by SN38+JQ1 treatment may proceed across replication boundaries, thereby interfering with replication origin firing and fork progression. We divided the genome into early and late replicating regions based on the Repli-seq signal using the RepliScan algorithm *(61)*. We found 15% of DoG transcripts were in late replicating regions while 85% of DoG transcripts were in early replicating regions. Notably, a few of the latter DoG transcripts were upstream of replication boundaries, with 8.2% (55 genes) within 200 kb of the early-to-late regions. Considering that SN38+JQ1 caused DoG transcription 200 kb beyond the TES (fig. S3E), this suggests that readthrough transcription can continue through replication boundaries. Indeed, the distance between DoG genes and early-to-late replication boundaries was markedly smaller than seen with non-DoG genes (Fig. 4D) and the direction of readthrough transcription typically travels from early to later replication regions (fig. S4B).

If readthrough transcription occurs in heterochromatic late replicating regions or at least in regions where transcription is typically repressed, it might affect chromatin compaction, leading to a de-repressed chromatin state. Indeed, the levels of H3K36me3 in late regions were reduced when comparing untreated and SN38+JQ1 treated samples (Fig. 4E). This change in chromatin status could in turn provoke activation of normally dormant origins, which usually do not fire under untreated conditions, as it is more likely that the replication fork originating from a more active origin reaches them beforehand. However, in conditions where replication is globally affected (e.g., SN38 treatment), dormant origins might get activated as previously seen *(62)*. We quantified the E and L Repli-seq reads in the early and late replicating regions (Fig. 4C, pink and blue boxes), respectively. In early replicating regions, both SN38 and SN38+JQ1 treatment showed an equal reduction in DNA replication suggesting that SN38+JQ1 treatment had no additional effect on early replication (Fig. 4F). In stark contrast, SN38+JQ1 treatment partially rescued replication in late replicating regions as increased signal could be measured relative to SN38 alone (Fig. 4G, compare red and green curves). Thus, the data supports our hypothesis that SN38+JQ1-driven readthrough transcription induces dormant origin firing in late replicated areas, specifically in conditions when early replication elongation is impaired by SN38.

### SN38+JQ1 treatment induces DNA damage in S-phase and cell stress signaling in G1 and G2 phases

TOP1 activity is required to resolve replication dependent supercoiling during S-phase *(28)*. If TOP1 is trapped on the DNA by TOP1 inhibitors, these can be converted to DNA double-stranded breaks (DSBs) in S phase by replication run-off *(63, 64)*. Furthermore, loss of TOP1 activity has been shown to inhibit replication fork progression and to induce replication-transcription interference likely due to supercoiling accumulation *(60, 65–67)*. This in turn can lead to cell cycle arrest and cell death *(68, 69)*. We predicted that SN38+JQ1-induced transcription into gene-free regions and late replication origin firing would increase the probability of replication fork stalling and DNA damage compared to TOP1 inhibition by SN38 alone.

To investigate this potential mechanism of drug synergy, we assessed the extent of DNA damage by quantification of DNA damage marker γH2AX *(70)* in S phase cells labelled by EdU incorporation. In agreement with previous research *(28)*, γH2AX was specifically upregulated in EdU+ S-phase cells in response to SN38 exposure (Fig. 5A, compare lane 1 *vs*. lane 2; and fig. S5A). This signal was significantly increased after SN38+JQ1 treatment indicating a synergistic DNA damage response (Fig. 5A, compare lane 2 *vs*. lane 4). We conceived two independent methods to test whether the DNA damage upon SN38+JQ1 was dependent on transcription interference with the replisome. The CDK9 inhibitor flavopiridol inhibits RNAPII elongation and can block both genomic and readthrough transcription *(71)* in asynchronous Bo103 cells (fig. S5B). On the other hand, the CDC7 inhibitor XL-413 is able to inhibit DNA replication initiation *(72)*, as shown by reduced EdU incorporation into replicating DNA independent of SN38+JQ1 treatment (Fig. 5B, compare lanes 5-8 *vs*. lanes 1-4). If the increased γH2AX seen upon SN38+JQ1 treatment stems from the transcription-dependent replication stress, co-treatment with either flavopiridol or XL-413 should revert SN38+JQ1-induced γH2AX signaling back to the level of SN38 treatment alone. Although flavopiridol treatment with SN38 alone increased γH2AX, in line with previous reports of flavopiridol potentiating the apoptotic effect of TOP1 poisons *(73)*, co-treatment with either flavopiridol (Fig. 5A, compare lane 6 *vs*. lane 8) or XL-413 (Fig. 5A, compare lane 10 *vs*. lane 12) reverted SN38+JQ1-induced γH2AX back to SN38 levels, demonstrating these treatments only abrogated the SN38+JQ1-specific effects.

**Fig. 5.**
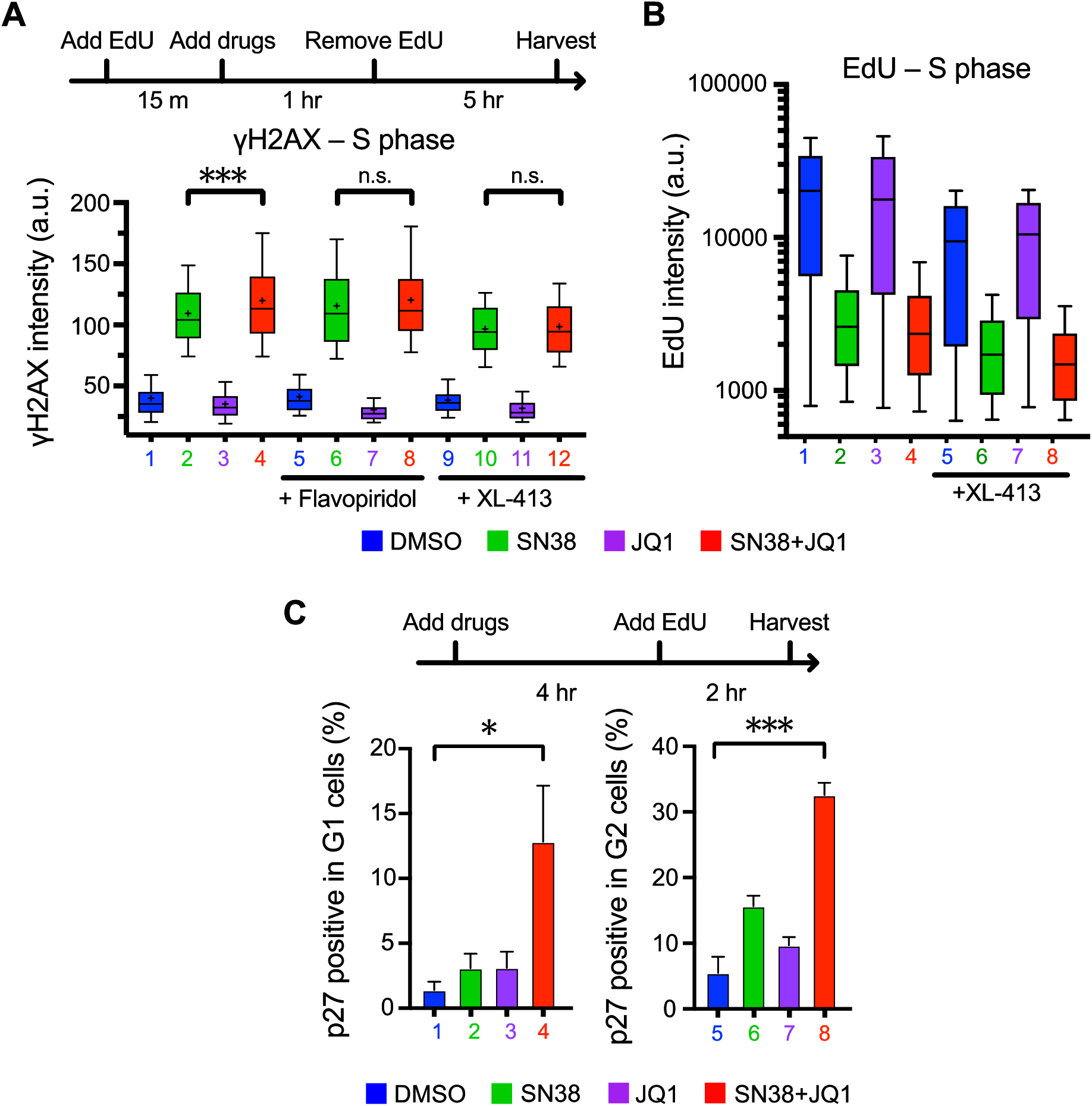
SN38+JQ1 induces readthrough transcription dependent replication stress in S-phase and cell stress signaling in G1 and G2 phases. **(A)** Top. Schematic of treatment. Bottom. Immunofluorescence quantitation of γH2AX intensity in S-phase (EdU positive) upon DMSO, SN38, JQ1, or SN38+JQ1 treatment +/-2 μM flavopiridol or 15 μM XL-413 for 6 hr. Mean represented by +, whiskers extend to 10-90%, a.u. = arbitrary units. Representative plot of n=3. ***: p<0.001; n.s.: not significant; Student’s t-test. **(B)** Flow cytometry analysis of EdU incorporation in S-phase cells after DMSO, SN38, JQ1, or SN38+JQ1 treatment +/-15 μM XL-413 after 6 hr. Whiskers extend to 10-90%, a.u. = arbitrary units. Representative plot of n=3. Top. Schematic of treatment. Bottom. Flow cytometry quantitation of p27 positive cells in G1 and G2 phase of the cell cycle upon DMSO, SN38, JQ1, or SN38+JQ1 treatment +/-2 μM flavopiridol after 6 hr (n=4, error bars represent standard deviation). *: p<0.05; ***: p<0.001, Student’s t-test.

We next tested whether the SN38+JQ1 treatment exhibited any synergistic effects in G1 and G2 cell cycle phases. The tumor suppressor p27 can arrest the cell in G1 under stress conditions to prevent potentially genotoxic consequences of DNA replication *(74)*. It can also cause G2 arrest in response to DNA damage in S-phase to avoid mitotic catastrophe *(75, 76)*. Persistent p27-dependent cell cycle arrest is shown to induce apoptosis in many cancer types *(77)*. Therefore, we postulated that p27 may become upregulated in G1 and G2 in response to replication interference induced by SN38+JQ1 treatment. We observed a clear synergistic upregulation of p27 expression in both G1 (Fig. 5C, compare lane 1 *vs*. lane 4) and G2 (Fig. 5C, compare lane 5 *vs*. lane 8) phases only after exposure to SN38+JQ1. Overall, these data suggest that the SN38+JQ1-induced readthrough transcription provokes replication stress and triggers a downstream DNA damage and stress signaling response.

### PDXs remain sensitive to Irinotecan+JQ1 treatment upon multiple treatment cycles *in vivo*

Having established a mechanistic model for the synergism of SN38+JQ1 in killing tumor cells, we then sought to translate our *in vitro* finding into preclinical settings to determine if we observe the same phenotypes *in vivo*. We performed RNA-Seq on the Bo99 PDX tumors described in fig. S1A, treated with Irinotecan or JQ1, alone or in combination. Analysis of the data showed a down-regulation of long genes after Irinotecan+JQ1 treatment (Fig. 6A and table S3), due of transcription inhibition. This effect was considerably more profound relative to the individual treatments, in contrast to the *in vitro* results (fig. S6A), suggesting that there may be even stronger synergy *in vivo*.

**Fig. 6.**
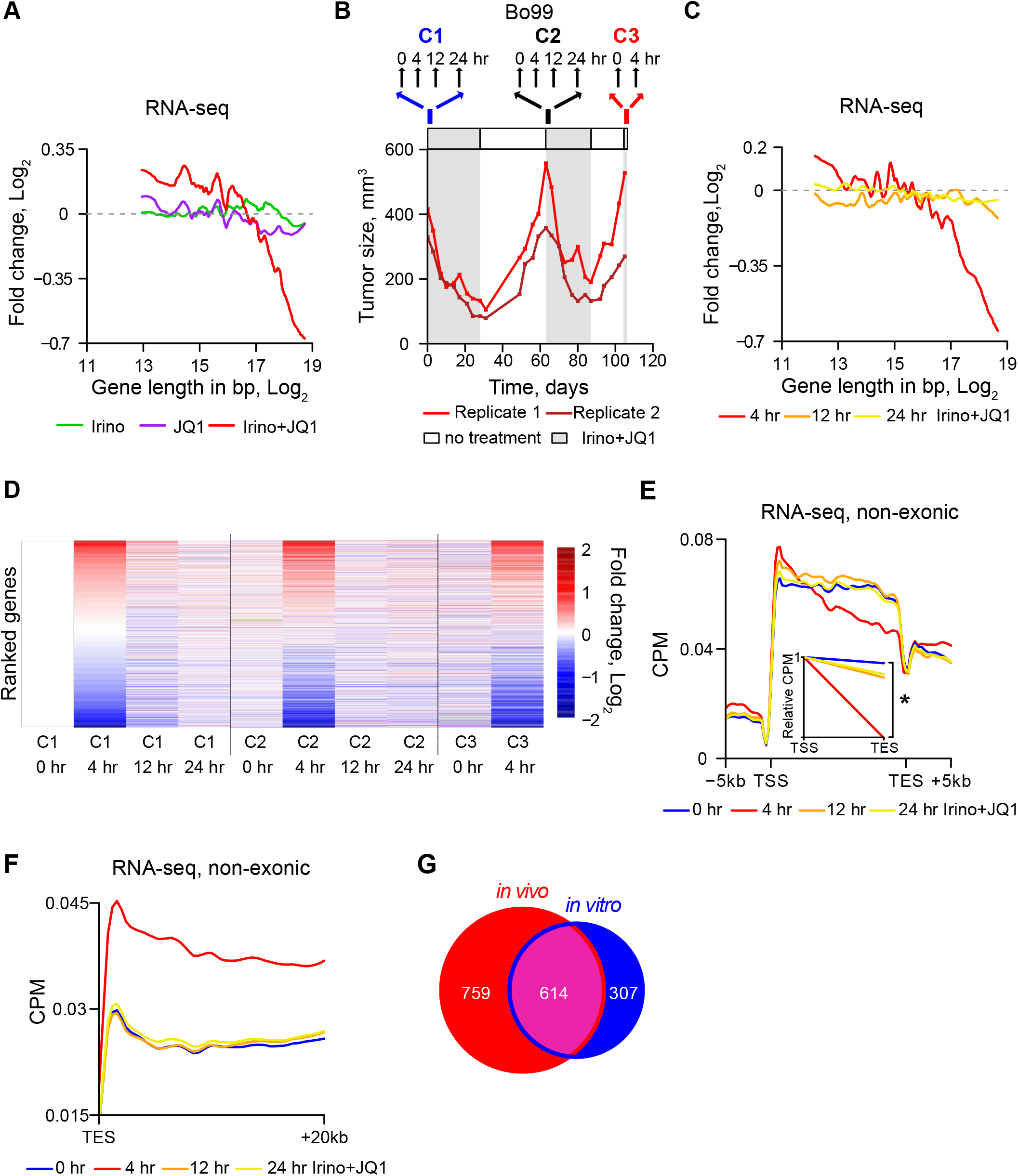
Irinotecan+JQ1 in patient-derived xenografts (PDX) triggers readthrough transcription and does not show emergent resistance over time. **(A)** Moving average of fold-change (Log_2_) derived from exonic RNA-Seq (hereafter indicated as RNA-seq) reads of treated (Irinotecan, JQ1, Irinotecan+JQ1) *vs*. untreated Bo99 PDX plotted against the gene length (Log_2_). Average of biological duplicates. **(B)** Top. Dosing and harvesting schedule. Bottom. Example growth curve of two Bo99 PDX tumors, subjected to 3 cycles of treatment with Irinotecan+JQ1. **(C)** Same as (A), but Bo103 PDX were treated with Irinotecan+JQ1 for 4, 12 and 24 hr and compared to untreated Bo103 PDX. Average of 3-4 tumors per condition. (C) Fold-change (Log_2_) of RNA-Seq reads of Bo103 PDX for each time point *vs*. untreated. Drug cycles 1, 2, and 3 = C1, C2, and C3. (**E)** Non-exonic RNA-Seq reads of Bo103 PDX plotted between the TSS and TES of the 10% longest protein-coding genes. Inset shows the gradient of the linear regression between TSS and TES (NERD index). Average of 3-4 tumors per condition. *: p<0.05, Student’s t-test. **(F)** Non-exonic RNA-Seq reads of Bo103 PDX plotted in the region 20 kb downstream of the TES of protein-coding genes. Average of 3-4 tumors per condition. **(G)** Venn diagram of DoG transcript producing genes detected in cultured Bo103 cells and in the Bo103 PDX after 4 hr of SN38/Irinotecan+JQ1 treatment.

To assess the early effects of Irinotecan+JQ1 on transcription, we designed an RNA-Seq protocol for Bo99 and Bo103 tissue material, performing the step of retro-transcription with random primers. This allowed detection of non-exonic reads and extrapolate information about nascent RNAs. We also devised a new treatment scheme in which we administered the PDX carrying mice with multiple cycles of Irinotecan+JQ1 (referred to as cycle 1, C1; cycle 2, C2; and cycle 3, C3), harvesting tumors 0, 4, 12 and 24 hr after the start of each treatment cycle (Fig. 6B). This way also allowed us to investigate whether tumor cells treated with the Irinotecan+JQ1 would develop resistance over time (tables S4 and S5). As expected, long genes were strongly downregulated in response to 4 hr treatment with Irinotecan+JQ1 (fig. S6B) and the tumors regressed upon subsequent treatment cycles (Fig. 6B and fig. S6, C to E), indicating that the cells do not gain resistance to the treatment. Common markers of resistance to either Irinotecan or JQ1 were not significantly dysregulated in the C3 0 hr timepoints relative to C1 0 hr, in agreement with continued drug sensitivity of the tumors to Irinotecan+JQ1 (fig. S6, F and G, and table S6). Long gene inhibition was lost 12 and 24 hr after treatment (Fig. 6C), indicating that the transcription inhibition was transient, and probably vanished as the drugs were metabolized and excreted. Overall, we found high concordance of the response after each treatment cycle and a general reversion after 12 hr (Fig. 6D).

By plotting the non-exonic reads onto the metagene, we then analyzed how the Irinotecan+JQ1 treatment affected transcription. First, we observed that 4 hr combination therapy provokes a buildup of short transcripts around the start of the genes, particularly of long genes, as seen *in vitro* (Fig. 2D). This accumulation dissipated after 12 and 24 hr (Fig. 6E and fig. S6H). Importantly, readthrough transcription was evident 4 hr after Irinotecan+JQ1 exposure (Fig. 6F), was found to extend as far as 500 kbs (fig. S6I) and was detectable at a highly similar set of genes as seen *in vitro* (Fig. 6G) with 45% of DoG genes from the *in vivo* experiment also detected *in vitro*.

### Irinotecan+JQ1 combination specifically targets transcriptionally-addicted tumors over normal cells

Tumor cells are dependent on transcriptional dysregulation to drive oncogenic growth *(2)*. This “addiction” to oncogenic drivers of gene expression can render tumor cells susceptible to disruption of transcription. If normal non-cancerous cells are less sensitive to transcription inhibition, such treatment would be more selective towards cancer cells and may provide a therapeutic window for clinical intervention. Therefore, if TOP1 and BRD4 inhibition treatment has clinical potential as a transcription targeting regimen, we would expect it to elicit a substantially reduced response in normal tissues, as we observed *in vitro* with normal immortalized pancreatic hTERT-HPNE cells (fig. S1F).

Since a subset of infiltrating mouse cells were co-harvested with each PDX tumor (6-12% in Bo99, 23-38% in Bo103), we could map RNA-Seq reads to the mouse genome to understand how Irinotecan+JQ1 treatment affected the expression of nascent RNAs in the mouse normal cells (tables S4 and S5). Interestingly, transcription of long genes was also reduced in mouse normal cells after 4 hr of Irinotecan+JQ1 but reverted after 12 and 24 hr, as seen in the human tumor cells (Fig. 7A and fig. S7A) indicating that transcription is targeted in both cell types. However, in contrast to the PDX cancer cells (Fig. 6E and fig. S6H), the NERD plot of the normal cells showed no substantial changes in the distribution of reads across the gene body after the different treatments (4 hr), either when observing all genes (fig. S7B) or long genes sensitive to elongation inhibition (Fig. 7B). Together, these data imply that while transcription elongation is inhibited also in the normal cells, the absence of short transcripts accumulating immediately downstream of the TSS suggests transcription does not continue to be initiated and subsequently stalled by trapped TOP1. Most strikingly, readthrough transcription was barely detectable in the non-malignant relative to the malignant cells (Fig. 7C).

**Fig. 7.**
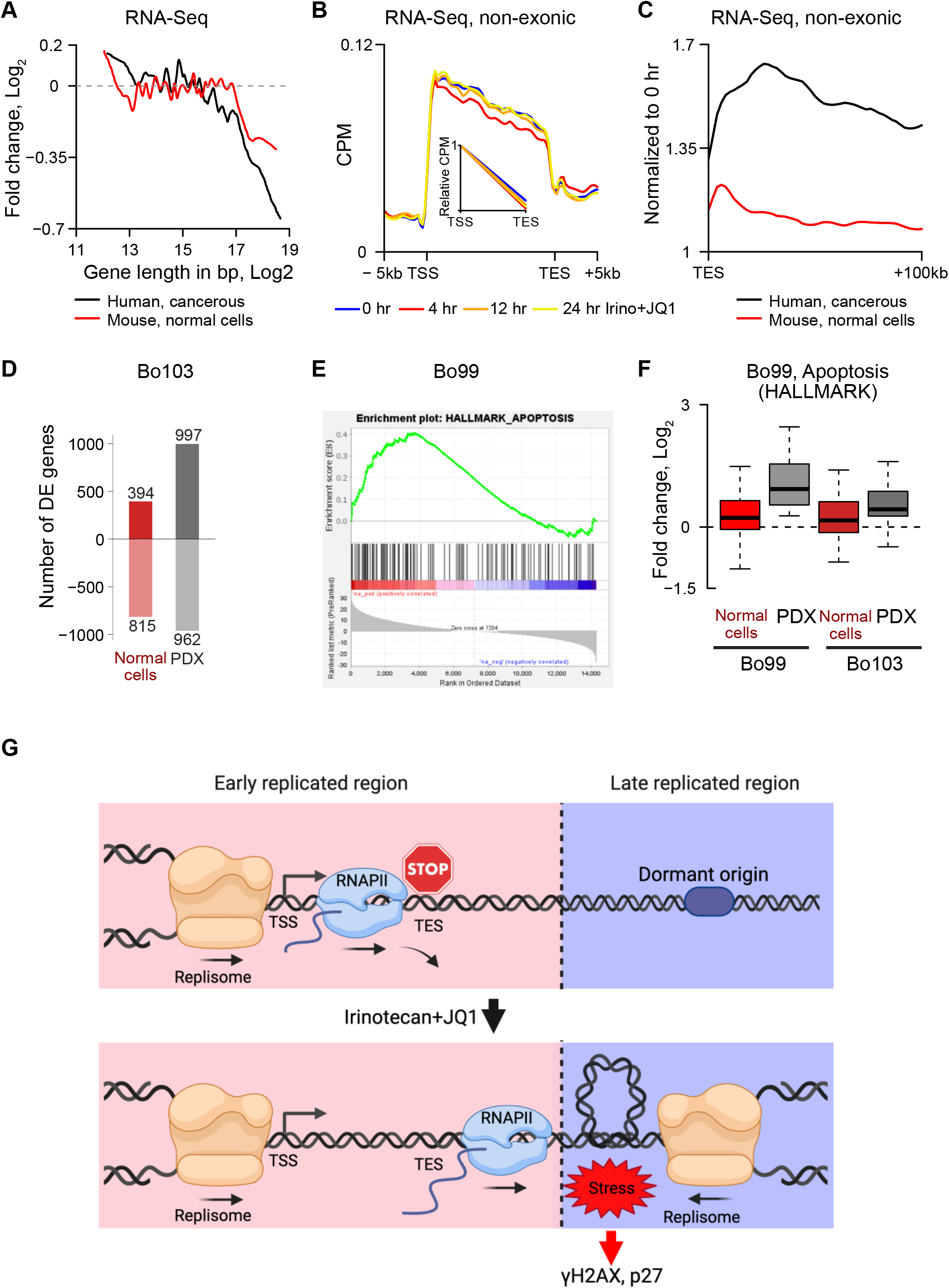
Transcription is preferentially affected in Bo103 PDXs upon Irinotecan+JQ1 compared to normal mouse cells. **(A)** Moving average of fold-change (Log_2_) of RNA-Seq reads from Bo103 PDX treated with Irinotecan+JQ1 for 4 hr and normal mouse cells. Fold change is plotted against the gene length. Average of 4 tumors per condition. **(B)** Non-exonic RNA-Seq reads from normal mouse cells plotted between the TSS and TES of the 10% longest protein-coding genes. Samples are isolated from Bo103 PDXs treated with Irinotecan+JQ1. Inset shows the gradient of the linear regression in between TSS and TES (NERD index). Average of 3-4 tumors per condition. No significant difference between NERD indexes is detected. **(C)** Non-exonic RNA-Seq reads of PDX tumors and normal mouse cells from Bo103 PDXs treated for 4 hr with Irinotecan+JQ1. Reads are plotted in the region 100 kb downstream of the TES of protein-coding genes. Data are expressed as counts per million (CPM) and normalized to the corresponding CPM values at 0 hr. Average of 4 tumors per condition. **(D)** Bar plot representing the number of statistically significant (adjusted p-value < 0.05) differentially expressed up- and down-regulated genes in Bo103 PDX tumor and associated mouse normal cells upon treatment with Irinotecan+JQ1 for 4 hr. **(E)** Gene Set Enrichment Analysis (GSEA) for the gene ontology term “Apoptosis” (NES=1.50, FDR=0.026) plotting gene enrichment after 4 hr Irinotecan+JQ1 treatment of Bo99 PDX tumor. **(F)** Boxplots representing the relative expression of the core enriched “Apoptosis” genes from (E) in Bo99 and Bo103 PDX tumors, both human cancer and mouse normal cells. **(G)** Working Model. Under untreated conditions, transcription and replication are coordinated with replication initiating in open, highly transcribed regions. Upon SN38+JQ1 treatment, readthrough transcription affect chromatin state and induce dormant origin firing in late replicated areas. This will interfere with the established replication timing pattern leading to replication stress, DNA damage and cell cycle arrest.

That the normal mouse tissue was not affected in promoter pausing regulation by Irinotecan+JQ1, indicated that 1) there should be fewer differentially expressed genes relative to the PDX tissues treated with the same drugs, and 2) signaling pathways involved in cellular stress should not be activated upon treatment. As expected, there were notably fewer differentially expressed genes in the normal compared to the PDX tissue (Fig. 7D and fig. S7C). While the number of down-regulated genes in PDX and normal samples are more similar in the Bo103 PDX due to inhibition of long genes (Fig. 7A), the number of upregulated genes was significantly greater in the PDX tissue indicating a stronger response to the treatment. Furthermore, gene set enrichment analysis (GSEA) *(78)* revealed that gene ontologies associated with cell death were not enriched in the mouse normal cells. The core genes from the hallmark gene ontology term “Apoptosis” were consistently upregulated in both Bo99 and Bo103 PDXs, but the homologous mouse genes in co-harvested normal mouse cells were not affected (Fig. 7, E to F).

All together, these results suggest that targeting transcription *via* dual inhibition of TOP1 and BRD4 can specifically target KRAS and MYC-driven pancreatic tumors and provide viable and clinically applicable therapy for hard-to-treat cancer entities such as PDAC and panNEC.

## DISCUSSION

While there have been recent advances made towards directly targeting KRAS^G12^ mutant tumors *(79, 80)*, selective targeting of oncogenic RAS activity is challenging and early clinical trials have been short lived *(81, 82)*. Therefore, many strategies are directed to target KRAS downstream signaling to indirectly block the oncoprotein’s effect *(83)*. Here we show that combining BET inhibition with Irinotecan to indirectly target oncogenic signaling is able to induce tumor regression, with two of the three PDAC PDX models tested reaching partial response, including one model of the difficult to treat basal (or quasi mesenchymal) subtype. Importantly the panNEC PDX model responded surprisingly well with complete response, suggesting that this new combination could be highly effective for both types of pancreatic cancer to be explored in future clinical studies. Considering the robust inhibition of KRAS-driven pancreatic tumors reported here, we propose that directly targeting transcription should be considered as a promising treatment strategy in clinical research.

SN38+JQ1 treatment promotes transcription downstream of the termination site by making the RNAPII not competent for interaction with TTFs downstream of the pausing site. Acute degradation of BRD4 has previously been shown to prevent TTF recruitment and efficient transcription termination, though JQ1 treatment alone was insufficient to cause extensive readthrough *(20)*. Interestingly, although JQ1 and SN38+JQ1 had similar effects on BRD4 displacement on chromatin (Fig. 2B), only the combination treatment caused significant loss of TTF recruitment and elevated readthrough transcription (Fig. 2, G to H) suggesting that loss of BRD4 binding cannot completely account for the readthrough transcription phenotype. The mechanism by which co-inhibition of BRD4 and TOP1 can synergistically prevent TTF recruitment remains unclear, particularly since the individual treatments showed a mild increase in TTF Cstf64 at the TSS (Fig. 2F). One possibility may involve the independent effects of both drugs on RNAPII phosphorylation. TOP1 poisoning by Camptothecin is known to release PTEF-b from its inactive complex with 7SK snRNP resulting in RNAPII hyperphosphorylation *(49)*. Additionally, in the absence of BRD4, RNAPII can be hyperphosphorylated through PTEF-b by the Super Elongating Complex (SEC) *(50, 85)*. The SN38+JQ1 treatment therefore would be expected to induce RNAPII hyperphosphorylation *via* two separate mechanisms: the release and activation of stored PTEF-b and the phosphorylation via SEC rather than BRD4. Indeed, our RNAPII-Ser2 ChIP-Rx-Seq data (Fig. 2F) highlight a JQ1-dependent increase in Ser-2 phosphorylated RNAPII at the TSS and a SN38-dependent increase towards the 5’ of the gene, with the SN38+JQ1 combination exhibiting both phenotypes. It is possible that the hyperphosphorylated state of RNAPII, along with the switch towards SEC-dependent activation preventing BRD4-dependent recruitment of TTFs, drives readthrough transcription. Indeed, the profound effects of flavopiridol, a potent inhibitor of PTEF-b *(86)*, on readthrough transcription strongly suggests that the observed effect is dependent on P-TEFb activation.

By demonstrating that readthrough is indeed sufficient to induce stress and causes both cell cycle arrest and apoptosis in transcriptionally addicted cancer cells (Fig. 7G), this study reveals the potential of inducing readthrough transcription to treat certain types of cancers. The differential response we present here between highly malignant human cancer cells and normal mouse cells suggest that targeting transcription and inducing readthrough transcription by BRD4+TOP1 co-inhibition may be particularly effective in transcriptionally addicted malignant cells. It thus represents a promising and clinically feasible treatment option, not only for the hard-to-treat PDAC and panNEC, but potentially also for other solid tumor types.

Cancer cells require high levels of transcription and replication to maintain oncogenic proliferation. Cell transformation can induce replication stress, subsequently leading to DNA damage, senescence, and cell death *(87, 88)*. Thus, to continue proliferating, cancer cells must adapt to tightly regulate the coordination between the replication and transcription machineries. This makes tumor cells susceptible to targeting replication stress therapeutically *(89)*. The importance of TOP1 and BRD4 to maintain optimal transcription is well established. However, both proteins are also involved in regulation of DNA replication. TOP1 is essential for removing supercoils that accumulate in response of replisome translocation *(90)*, while BRD4 regulates DNA replication checkpoint signaling *(91)*. Therefore, characterization of the synergistic effect of TOP1+BRD4 co-targeting must account for the independent roles of replication and transcription, and how these processes interact. We show that replication stress in response to SN38+JQ1 treatment is dependent on both dysregulated transcription and replication dormant origin firing (Fig. 5A), highlighting the dual targeting of this combination therapy.

There has been a resurgence of interest in readthrough transcription over the last ten years *(92)*. This process can be triggered by heat or osmotic shock, or by viral infection *(71, 93)*. It has been described to be driven by loss of termination factors, as we demonstrate in our study, or by disruption of other RNA processing factors such as Integrator *(94)*. The purposes and outcomes of readthrough transcription are still being elucidated. Some reports place this process downstream of stress signaling to protect the cell from external stresses. For example, the long ncRNA produced by readthrough transcription can act as a nuclear scaffold to maintain nuclear integrity after osmotic stress *(95)*. However, in other instances such as during Influenza A Virus (IAV) infection, readthrough transcription can directly generate a stress response by remodeling the 3D structure of the genome *(71)*. In this regard, the IAV protein NS1 induces transcriptional readthrough, which continues through topologically associated domain (TAD) boundaries. This displaces cohesin from TAD boundaries and decompacts the heterochromatic DNA region to create a more permissive chromatin *(71)*. Here, we demonstrate that readthrough transcription driven by SN38+JQ1 treatment is frequently proximal to replication boundaries, which are largely congruous with TADs *(96)*, and exhibits many phenotypes consistent with permissive heterochromatin including loss of the silencing histone marker H3K36me3 and dormant origin firing in late S-phase replication. Interestingly, IAV infection has also been demonstrated to upregulate the stress response protein p27 *(97)*, supporting our data (Fig. 5C). Interestingly, the elevated replication in heterochromatic compartments was not strictly isolated to regions of readthrough transcription suggesting that local chromatin decompaction at DoG transcription sites can lead to broad decondensation of late replicating heterochromatin. Recent research into heterochromatin maintenance through phase separation *(98, 99)* may suggest that readthrough transcription could disrupt the entire phase-separated compartment, thus propagating chromatin compartment switching at a distance.

Further evidence supporting the concept that the SN38+JQ1 response phenotype is related to aberrant transcription can be extrapolated from our earlier work. We previously showed that the colon carcinoma HCT116KI cell line, which has an exon 4 deletion in TOP1 preventing RNAPII-TOP1 interaction, without affecting the enzymatic activity of TOP1, exhibited greatly reduced synergy between SN38 and JQ1 compared to the isogenic wild type cell line *(16)*. While these cancer cells showed clear perturbation in transcription through loss of coordination between transcription and supercoil relief, they are equally sensitive as wild type cells to inhibition of replication by fluorouracil, suggesting that replication is not impaired in the HCT116KI cells *(100)*.

Another potential advantage of directly targeting oncogenic transcription is the likely reduced chance of resistance development upon repeated treatment cycles. Precision medicine, the therapeutic targeting of specific drivers of cancer, can initially elicit rapid tumor regression, but sub-populations of tumors emerge driven by alternative pathways which are resistant to continued therapy *(101)*. However, in contrast to the redundancy of signaling pathways, transcription is an essential and irreplaceable feature of cellular homeostasis. Our work indicates that the downstream response to the drugs with regards to readthrough transcription and stress signaling is profoundly elevated in the PDX-containing malignant cells relative to the normal mouse cells from the same lesion (Fig. 7). Since cancer cells are “addicted” to elevated transcription rates, the direct targeting of oncogenic transcription provides a therapeutic window for precision treatment.

This study was limited to characterizing the response to TOP1 and BRD4 inhibition both *in vitro* by cell culture and *in vivo* with PDX models. Neither of these systems take into consideration the potential effects of a functional immune system on drug response, with respect to both tumor killing and tolerance of the host. Further studies using genetically engineering mouse models of PDAC will be essential to judge the clinical suitability of this therapeutic strategy. On this point, the use of BET inhibitors in the clinical setting is somewhat restricted due to the observed dose limiting toxicities *(84)*. However, we saw synergy across a range of JQ1 doses (Fig. 1G), suggesting this combination may allow for lower dosing of BET inhibitors. It will be interesting to address the selective role of the BD1 and BD2 domains of BRD4 here further inaugurating potential strategies with lower toxicity and thus clinical applicability. Finally, while we demonstrated that the SN38+JQ1 DNA damage was depended on transcription using flavopiridol (Fig. 5), we were unable to specifically inhibit readthrough transcription without also blocking transcription in general. Therefore, we cannot completely exclude the possibility that some unknown transcription-dependent feature of SN38+JQ1 treatment underlies the DNA damage response. As the readthrough transcription phenomenon continues to be elucidated by us and others, we hope to acquire the tools to further investigate the mechanism of action of combined BRD4 and TOP1 inhibition.

## MATERIALS AND METHODS

### Study design

The objective of this study was to test co-inhibition of TOP1 and BRD4, which we previously demonstrated to be synergistic *in vitro (16)*, in PDX models of pancreatic carcinoma to determine whether this treatment strategy could be effective *in vivo*. We hypothesized that targeting transcription through two independent arms would selectively kill tumor cells that are oncogenically addicted to transcription, while leaving normal cells unharmed. We also predicted that by targeting a fundamental feature of cancer proliferation (as opposed to a specific driver of oncogenesis), we would avoid the emergence of drug resistance often associated with targeted therapy. For this, we used a selection of PDAC and panNEC PDX models either treated acutely (4-24 hr) to assess response to treatment, or repeatedly, stopping drug administration once the tumor has regressed and recommencing upon tumor progression, to establish whether the tumors remain sensitive to treatment. All transplanted mice were randomized into treatment groups. Mouse experiment endpoints were defined by the associated ethics approval to safeguard the health of the mice. The Bo103 PDAC was adapted for tissue culture to enable investigation of the mechanism of action *in vitro*. No data was excluded from these studies. Further information about the study design, number of replicates per experiment and measurement methods can be found in the Supplementary Materials and Methods.

### Statistical analysis

All statistical tests were performed using Graphpad Prism version 8 or 9 or statistical functions in R using the tests described for each experiment. *: p < 0.05; **: p < 0.01; ***: p < 0.001. Information about statistical tests is provided in figure legends for respective figures and in the Supplementary Materials and Methods.

## List of Supplementary Materials

Materials and Methods

Figs. S1 to S6

Tables S1 to S6

References *(103-124)*

## Acknowledgements

We would like to thank Arne Lindqvist, Fedor Kouzine, Bennie Lemmens, Camilla Björkegren and David Levens for critical review of the manuscript. We thank Christian Sommerauer for providing technical expertise with the sequencing machine, and Margaret Sällberg Chen for sharing the hTERT-HPNE cell line. The computations and data storage were enabled by resources in project [SNIC 2018/8-390], provided by the Swedish National Infrastructure for Computing (SNIC) at UPPMAX, partially funded by the Swedish Research Council through grant agreement no. 2018-05973. This work was supported by the Karolinska Institutet Biomedicum Flow Cytometry and Imaging Core Facilities.

## Funding

Knut och Alice Wallenbergs Stiftelse KAW 2016.0161 (LB)

Vetenskapsrådet 2016-02610 VR (LB)

Vetenskapsrådet 2021-02630 VR (LB)

VINNOVA 2016-02055 (LB)

Cancerfonden CAN 2018/760 (LB)

Cancerfonden Pj 21 1771 (LB)

Karolinska Institutet KID Grant 2018-02013 (LB)

Georgius Agricola Stiftung Ruhr, Bochum (SAH)

Deutsches Konsortium für Translationale Krebsforschung (JTS)

Deutsche Forschungsgemeinschaft 405344257/SI1549/3-2 (JTS)

Deutsche Forschungsgemeinschaft SI1549/4-1 (JTS)

Deutsche Krebshilfe #70112505/PIPAC (JTS)

Bundesministerium für Bildung und Forschung 01KD2206A/SATURN3 (JTS)

## Author Contributions

Conceptualization: DPC, JTS, SAH, LB

Software: JG, VK, EI, PFC

Investigation: DPC, JG, SwL, VK, EI, AW, HB, STL, SmL, PFC, DV, SAH

Resources and Data Curation: MP, RV, CT, HW, SU, AT, WS, KG, CK, LB

Writing – Original Draft: DPC, JG, VK, EI, AW, MA-H, SAH, LB

Writing – Review and Editing, All authors

Supervision: MA-H, SAH, LB

Funding Acquisition: JTS, SAH, LB

## Data and materials availability

Data relating to this paper will be released upon peer-reviewed publication on the Gene Expression Omnibus (Accession# GSE211471). Materials should be requested from Dr Stephan A. Hahn and Dr Laura Baranello.

