## Supplementary Methods and Figures for "Co-inhibition of topoisomerase 1 and BRD4-mediated pause release selectively kills pancreatic cancer *via* readthrough transcription"

#### Supplementary Materials and Methods

##### PDX Biobank

Establishment of the PDX mouse model was performed using surgically resected PDAC and panNEC tissues collected from patients at the Ruhr-University Bochum Comprehensive Cancer Center. Informed and written consent was obtained from all patients. The study was approved by the ethics committee of the Ruhr University Bochum (permission no. 3534-09, 3841-10, 16-5792). Patient tumor tissues were xenografted in both flanks of nude mice (Janvier, St Berthevin Cedex, France), expanded, isolated, and re-implanted for at least three generations. 5-10 week old mice were housed in specific pathogen free conditions where light, temperature (+ 21°C), and relative humidity (50-60%) were controlled. Food and water were available *ad libitum*. All animal experiments were approved by the local authorities (81-02.04.2017.A423) and performed in accordance with the guidelines for Ethical Conduct in the Care and Use of Animals. PDAC PDX class assignment was defined based on consensus clustering on median-centered data for the top 3000 most variable features after signal normalization. To this end, non-negative matrix factorization was used for discovery and validated by consensus clustering with the help of the consensus cluster plus algorithm for determining cluster count and membership by stability evidence in unsupervised analysis. The algorithm begins by subsampling a proportion of items and a proportion of features from a data matrix. Each subsample is then partitioned into up to k groups by a user-specified clustering algorithm: agglomerative hierarchical clustering, k-means or a custom algorithm. This process is repeated for a specified number of repetitions. Pairwise consensus values, defined as ‘the proportion of clustering runs in which two items are [grouped] together’ (103), are calculated and stored in a consensus matrix (CM) for each k. Then for each k, a final agglomerative hierarchical consensus clustering using distance of 1–consensus values is completed and pruned to k groups, which are called consensus clusters. The algorithm was benchmarked with the Collisson subtypes and they applied to the PDX data.

##### Treatment Cohorts

To establish treatment cohorts, tumor pieces (1-2 mm) from early passage PDXs (≤ F4 generation) were soaked in undiluted matrigel (Becton Dickinson) for 15 to 30 min. and subsequently implanted subcutaneously onto female mice (NMRI-Foxn1nu/Foxn1nu, Janvier, St Berthevin Cedex, France) at two sites (scapular region) using as many as 4 pieces per site. Tumors were allowed to grow to a size of approx. 100-200 mm<sup>3</sup>, at which time mice were randomized in the treatment and control groups with five to six mice in each group. Tumor volumes were estimated from 2-dimensional tumor measurements by caliper measurements twice a week using the following formula: Tumor volume (mm<sup>3</sup>) = $(\pi/6)(D)(d^2)$, where d is the minor tumor axis and D is the major tumor axis. Mice were treated daily with JQ1 (Hycultec) 50mg/kg by i.p. injection, and with Irinotecan (Hycultec) i.p. three times per week (Mon., Wed., Fri.) at 15mg/kg, followed by one week of treatment pause for two consecutive cycles. OTX015 (Hycultec) was given at 50mg/kg i.p. for five consecutive days followed by two days of treatment pause.

##### Multiplexed IF (mIF) histological staining

Multiplexed IF was performed using the Opal multiplex system (Perkin Elmer, MA) according to the manufacturer's instruction and as previously described (104). In brief, FFPE sections were deparaffinized and then fixed with 4% paraformaldehyde prior to antigen retrieval by heat-induced epitope retrieval using citrate buffer (pH6) or Tris/EDTA (pH9). Each section was put through several sequential rounds of staining; each includes endogenous peroxidase blocking and non-specific protein blocking, followed by primary Ab and corresponding secondary horseradish peroxidase-conjugated polymer (Zytomed Systems or Perkin Elmer). Each horseradish peroxidase-conjugated polymer mediated the covalent binding of different fluorophores using tyramide signal amplification. Such covalent reaction was followed by additional antigen retrieval in heated citrate buffer (pH6) or Tris/EDTA (pH9) for 10 min to remove antibodies before the next round of staining. After all sequential staining reactions, sections were counterstained with DAPI (Vector lab). The sequential multiplexed staining protocol used primary antibodies for cleaved caspase 3, γH2AX and PanCK, and with secondary antibodies labelled with Opal 570, 480 and 520, respectively. Slides were scanned and digitalized by Zeiss AxioScanner Z.1 (Carl Zeiss AG) with 10x objective magnification. Quantification of individual and/or co-expressing markers in the mIF images was performed with HALO (Indica Labs).

#### Cell culture

To establish the primary cell line the corresponding Bo103 PDX tumor was harvested and subsequently dissociated by gentleMACS dissociator (Miltenyi Biotec). The human tumor dissociation kit (Miltenyi Biotec) and mouse cell depletion kit (both from Miltenyi Biotec) were utilized according to the manufacturer's protocol. Fibroblasts were removed by differential trypsinization. The identity of Bo103 was confirmed by comparing short tandem repeat (STR) data between the Bo103 PDX founder tumor and the derived primary cell line. STR analyses were performed as previously described (105) and analyzed on a CEQ8800 sequencer (Beckman Coulter). Bo103 cells were grown in a 1:1 mix of high glucose (4.5 g/L) DMEM (Thermo, 61965059) and DMEM/F12 (Thermo, 11320033) containing 5% FBS (Thermo, 10270106), 1.6 μg/ml amphotericin (Sigma, A2411), 10μM Y-27632 (Selleckchem, S1049), 6.2 μg/ml ciprofloxacin (Sigma, 17850), 8.4 ng/ml cholera toxin (Sigma, C8052), 500 ng/mL insulin (Sigma, I9278) and 20 nM 1-thioglycerol (Sigma, M6145), in a 37°C incubator supplied with 5% CO<sub>2</sub>. All *in vitro* drug treatments use 500 nM SN38 and 1 μM JQ1, unless otherwise noted. To enrich cells in S phase, cells were blocked in S phase by addition of 200 mM hydroxyurea for 18 hr. hTERT-HPNE cells were cultured in high glucose DMEM media containing 5% FBS, 2mM GlutaMAX (Thermo, 35050061), 10ng/ml EGF (Thermo, PHG0314) and 750ng/ml puromycin (Thermo, A11138-03), in a 37°C incubator supplied with 5% CO<sub>2</sub>.

#### Antibodies

Primary antibodies used in this study are anti-Top1 (Abcam, ab109374), anti-BRD4 (Thermo, A301985A), anti-H3K36me3 (Abcam, ab9050), anti-H3K27ac (Abcam, ab177178), anti-Cstf64 (Thermo, A301092A), anti-RNAPII-Ser2-P (Abcam, ab5095), anti-γH2AX for *in vitro* IF (Sigma, 05-636), anti-γH2AX for mIF (CST, 9718S), anti-p27 (Abcam, ab32034), anti-BrdU

(BD, 555627), anti-cleaved caspase 3 (CST, 9664L) and anti-PanCK (Abcam, ab6401). Secondary antibodies are anti-mouse (Thermo, A11029) and anti-rabbit (Thermo, A32790) Alexa Fluor 488 conjugated antibodies.

#### **Viability assay**

Bo103 or hTERT-HPNE cells were plated in 96-well plates at 4000 cells/well. After 48 hr, cells were treated with 0.01-30 μM of SN38 and JQ1 in a checkerboard format in a final volume of 200 μl. After a further 48 hr, media was replaced with 50 μl room temperature media and the plates were incubated at room temperature for 30 min. Then, 50 μl of Cell Titer-Glo 2.0 (Promega, G9242) was added to each well, the plates were placed on an orbital shaker for 2 min to lyse cells, then well luminescence was measured using a FLUOstar Omega Microplate Reader (BMG Labtech). The relative viability was calculated by normalizing signal to the viability of media-only wells. The Bliss coefficient was determined as the difference between the measured effect of the drugs, and the predicted effect if the drugs act independently. This predicted effect was calculated as $d_1 + d_2 - d_1d_2$, where $d_1$ and $d_2$ are the effect of each drug alone at the relevant concentrations.

#### **qPCR**

Total RNA was extracted from Bo103 cells, using the NucleoSpin RNA kit (Macherey-Nagel) according to the manufacturer's instructions. cDNA synthesis was performed with 1 μg RNA in a 2-stage reaction, first annealing random primers (Promega) at 65°C for 5 min and then ice for 1 min followed by reverse transcription using SuperScript<sup>TM</sup> IV Reverse Transcriptase (Invitrogen, Thermo Fisher Scientific) at room temperature for 10 min, following incubation at 50°C for 30 min, then inactivation at 80°C for 10 min. Readthrough transcription analysis was performed with primers sets (SERP1 – F: TCAGGACCTAGGTTTACTGAAGA, R: TCTTCTCTGCCCTAGCCCAA; SEC61B – F: CCTCCAGTTCTGGGTGGTTC, R: GACAGACAAGCCAGCAGCTA; BPNT2 – F: TATCACCACCCTGCCTTGTG, R: TTGGCATCCATGCTCCGATT) targeting specific regions approximately 60 kb downstream of the transcription end site of the target gene. qPCR reactions were carried out on a CFX96 Real-time system device (BioRad Laboratories) using the Fast SYBR<sup>TM</sup> Green Master Mix (Applied Biosystems, Thermo Fisher Scientific), according to the manufacturer's instructions. All reactions were run in duplicates in three independent experiments. The $\Delta\Delta C_t$ method was used for data analysis and fold changes in gene expression were normalized relative to the DMSO treated sample.

#### **Chromatin immunoprecipitation with reference exogenous genome (ChIP-Rx)**

H3K27ac, H3K36me3, RNAPII-Ser2-P, BRD4 and Cstf64 ChIP were performed on Bo103 cells as described previously (16) with minor modifications. Chromatin from mouse embryonic fibroblast (MEF) cells was used to spike-in the human PDX chromatin to enable normalization across samples. Briefly, Bo103 cells were treated for 4 hr with stated drug concentrations. Bo103 and MEF cells were crosslinked with 1% formaldehyde (Thermo Fisher, 28906) for 5 min. Cross-linking was stopped by the addition of glycine (125 mM) and cells were washed twice with cold PBS. After harvesting cells by scraping, the pellet was washed once with PBS

plus 0.5% BSA and resuspended in RIPA buffer (10 mM Tris-HCl pH 8.0, 1 mM EDTA pH 8.0, 1 % TritonX-100, 0.1 % Na-Deoxycholate, 0.1 % SDS, 200 mM NaCl, with the addition of protease inhibitor cocktail) to a final concentration of $1 \times 10^7$ cells/ml. Samples were sonicated with Bandelin probe and Covaris ME220 sonicators to produce chromatin fragments of 400 bp on average. After centrifugation, extracts were immunoprecipitated. 2 μg of anti-RNAPII, 3 μg of anti-BRD4, 5 μg of anti-Cstf64, 2 μg anti-H3K27ac or 1 μg anti-H3K36me3 were mixed with 35 ml Protein A/G magnetic beads (Pierce, 88803) and incubated at 4°C for 6 hr with controlled rotation. For ChIP-Rx, Bo103 chromatin from $3 \times 10^6$ Bo103 cells was mixed with MEF chromatin at a 5:1 ratio. The mixture was incubated with Protein A/G-antibody complexes with rotation overnight at 4°C. Samples were washed twice with RIPA buffer, twice with RIPA buffer containing 300 mM NaCl, twice with LiCl buffer (10 mM Tris-HCl pH 8.0, 1 mM EDTA pH 8.0, 250 mM LiCl, 0.5 % NP40, 0.5 % Na-Deoxycholate) and twice with TE (10 mM Tris-HCl pH 8.0, 1 mM EDTA pH 8.0). The beads were then resuspended in 125 ml TE plus 0.25 % SDS supplemented with 60 μg proteinase K (500 μg/ml, NEB, P8107S) and incubated overnight at 65°C. DNA was either recovered from the elute by phenol:chloroform:isoamyl alcohol (25:24:1) (Sigma, P2069) extraction followed by ethanol (100%) precipitation in the presence of 20 mg of GlycoBlue (Thermo Fisher, AM9515), or using the QIAquick PCR Purification Kit (Qiagen, 28106) and dissolved/eluted in Tris-HCl pH 8.5. All ChIP-Rx-Seq experiments were performed in biological duplicates.

###### **Covalent adduct detection coupled to ChIP-Seq for TOP1 (TOP1 CAD-Seq)**

Following Kuzin et al. (37), about $1 \times 10^7$ Bo103 cells were treated (in biological duplicates for the sequencing assays) with SN38 alone or in combination with JQ1 for 1 hr. During the last 30 min, MG132 (10 μM) was added to the cells. Cells were immediately lysed in 2 ml of M buffer (9.3 mM Tris-HCl pH 6.5, 18.6 mM EDTA, 5.59 M guanidine thiocyanate, 0.93 % DTT, 0.93 % Sarcosyl, 3.72 % TritonX-100) and briefly sonicated with Bandelin probe sonicator at 20 % amplitude for 3 cycles with 30 sec ON, 30 sec OFF. DNA covalent adducts were precipitated with 50% EtOH at -20°C, centrifuged at 14000 rpm and pellets were washed thrice in wash buffer (20 mM Tris-HCl pH 7.5, 50 mM NaCl, 1 mM EDTA, 50 % EtOH). Pellets were dried for 5 min and resuspended in TE-SDS 0.1 % (10 mM Tris-HCl pH 8.0, 1 mM EDTA pH 8.0, 0.1 % SDS). After 30 min incubation by gentle agitation samples were further sonicated with Covaris ME220 sonicator for 5 min at High Cell protocol in milliTUBE–1 mL with AFA Fiber to produce fragments of about 1 kb. For the immunoprecipitation, 2 μg anti-TOP1 were mixed with 30 μl Protein A/G magnetic beads (Pierce, 88803) and incubated at 40°C for 6 hr with rotation. Beads were washed once with ice-cold PBS and DNA covalent adducts from $1 \times 10^7$ cells were added to the Protein A/G-antibody complexes and incubated overnight at 40°C with rotation. Washing was performed as described for the ChIP protocol but only once with every buffer and always in presence of 0.1% SDS. The beads were then resuspended in 100 μl TE plus 0.5% SDS supplemented with proteinase K (500 μg/ml) and incubated for 4 hr at 65°C. Samples were then purified by QIAquick PCR purification Kit.

###### **FACS**

Bo103 cells were treated for 6 hr with 500 nM SN38, 1 μM JQ1, 2 μM flavopiridol and/or 15 μM XL-413. For the final 2 hr, cells were treated with 20 μM EdU to label replicating cells. Cells were harvested by trypsinization, washed in PBS, and fixed by dropwise addition of chilled 70% ethanol while vortexing, and stored at -20°C. Cells were washed twice in PBSTB (PBS, 0.1% TritonX-100, 1% BSA), then the EdU was conjugated to Alexa Fluor 647 via click chemistry using the Click-iT EdU Flow Cytometry Assay Kit (Invitrogen). The samples were washed in PBSTB and incubated with 1:1000 dilution of primary antibody at room temperature for 30 min, then washed and incubated with secondary anti-mouse or anti-rabbit antibody conjugated to Alexa Fluor 488 in the dark at room temperature for 30 min. Finally, cells were washed and incubated in PBSTB with 1:1000 FxCycle Violet and 10 μg/ml RNase for 15 min in the dark to stain for DNA content. Samples were assayed on the FACS Canto II (Becton Dickinson). Cells were categorized into cell cycle stages by DNA content and EdU positivity. All experiments were repeated 3-4 times.

##### **Microscopy**

Bo103 cells were plated in 96-well tissue culture plates at 8000 cells/well and grown for 48 hr. Cells were treated for 6 hr with 500nM SN38, 1 μM JQ1, 2 μM flavopiridol and/or 10-15 μM XL-413. Cells were pulsed 15 min before drug treatment with 20 μM EdU, which was washed out 1 hr after drug treatment and wells were replaced with drug-containing media. The media was aspirated, and cells were fixed with 3% paraformaldehyde/PBS at room temperature for 5 min before washing twice with PBS. Cells were permeabilized both by incubation at -20°C in methanol for 2 min and, subsequently after washing twice in PBS, by incubation in PBS + 0.5% TritonX-100 at room temperature for 10 min. The EdU was conjugated to Alexa Fluor 647 using the Click-iT EdU Flow Cytometry Assay Kit (Invitrogen), before the cells were blocked in PBSTB for 1 hr at room temperature. The wells were sequentially incubated for 1 hr with 1:250 primary γH2AX antibody and 1:1000 secondary Alexa Fluor 488 antibody with 1 μg/ml DAPI, with three PBSTB washed following each incubation. The plate was stored in PBS and imaged on the ImageXpress Micro cellular imaging system (Molecular Devices) and analysed using CellProfiler 4 (106). Cells in S phase were identified based on EdU positivity. The experiment was independently repeated three times, and in all three instances the γH2AX signal for SN38+JQ1 was significantly higher than SN38 alone, while co-incubation with either flavopiridol or XL-413 ablated significant differences.

##### **Repli-seq**

This method was adapted from a published protocol (59). Bo103 cells were treated in duplicate with vehicle, SN38, JQ1, or both drugs, then with 100 μM BrdU, 4 hr and 2 hr before harvest, respectively. Cells were trypsinized, resuspended in 2.5 ml cold FACS buffer (PBS + 1% FBS), fixed with dropwise addition of 7.5 ml ice-cold 100% ethanol while vortexing, and stored at -20°C. After washing twice in FACS buffer, $3 \times 10^6$ cells were stained for DNA content in 300 μl FACS buffer + 50 μg/ml propidium iodide + 250 μg/ml RNase. $40-120 \times 10^3$ cells were sorted from the early S and late S populations for each sample. Cells were lysed in SDS-PK buffer (50 mM Tris pH8, 10 mM EDTA, 1M NaCl, 0.5% SDS, 200 μg/ml Proteinase K) at 56°C for 2 hr, then DNA was extracted using the Zymo Quick-DNA Microprep Kit and eluted in 130 µl H<sub>2</sub>O. DNA was sonicated to fragments of 200-300 bp in microtubes with the Covaris ME220 sonicator using the following settings: Peak Incident Power – 70W, Duty Factor – 20%, Cycles/burst – 1000, Time – 130 sec. Each sample was ligated with adapters using the NEBNext Ultra II DNA Library Kit and the adapter hairpins were processed with the USER enzyme and purified with the DNA Clean and Concentrator Kit (Zymo, D4013). DNA was denatured to ssDNA by incubation at 95°C for 5 min, then on ice for 2 min. Each sample was diluted in IP buffer (10 mM sodium phosphate pH7, 140 mM NaCl, 0.05% TritonX-100) and incubated with 0.5 µg anti-BrdU for 20 min at room temperature while rocking. Protein A/G Magnetic Beads were added and followed by incubation for a further 30 min, before washing the beads twice with cold IP buffer. The DNA was eluted by incubation in digestion buffer (50 mM Tris pH8, 10 mM EDTA, 0.5% SDS) with 50 µg Proteinase K overnight in a 37°C air incubator, then by adding a further 25 µg Proteinase K and incubating at 56°C shaking for 1 hr. The eluted ssDNA was purified with the DNA Clean and Concentrator Kit (Zymo Research). All samples were amplified and indexed using the NEBNext Multiplex Oligos for Illumina, primer-dimers were removed by Ampure XP enrichment (Beckman Coulter) and sequenced on the Illumina NextSeq 550. The sequencing run was single end with 75 bp reads.

##### **SLAM-seq**

A modified version of SLAM-seq (46) was used to measure both, nascent RNAs and steady state RNAs, over the full transcript length. 1.5x10<sup>6</sup> Bo103 cells were treated with DMSO, 500 nM SN38, 1 µM JQ1, or SN38+JQ1 for 4 hr and in the last 2 hr 100 µM 4-Thiouridine (S4U) was added to the medium in the dark (in triplicates). Cells were scraped stepwise in 2x0.5 ml Trizol (Invitrogen, 15596018), snap frozen and stored at -80°C. For RNA isolation we followed the manufacturer's instructions of SLAM-seq Kinetics Kit – Anabolic Kinetics Module (Lexogen, 061.24) except for the following modifications: frozen samples were thawed for 5 min at 65°C and mixed with 200 µl chloroform:isoamylalcohol mix (24:1) before further incubation for 15 min at 65°C (strongly shaking and vortexing from time to time). RNA in the aqueous phase was separated by centrifugation, supplemented with reducing agent and again extracted with chloroform:isoamylalcohol mix (24:1) before precipitation overnight at -20°C (in the presence of reducing agent). RNA was washed twice with 1 ml and 0.18 ml 75% EtOH (plus reducing agent), respectively and pelleted for 10 min at 7500g. Resuspension in H<sub>2</sub>O was facilitated by incubation for 15 min at 55°C and 10 min on ice and repeated pipetting. RNA was quantified with the Qubit RNA nano Assay Kit (Thermo Fisher, Q33230). 4.5 µg of the RNA was alkylated by addition of Iodoacetamide following the manufacturer's instructions. Qubit RNA nano Assay Kit (Thermo Fisher, Q33230) was used to quantify the RNA and RNA integrity was controlled using the RNA 6000 nano kit (Agilent, 5067-1511) on a Bioanalyzer 2100 (Agilent). Ribosomal RNAs were depleted from 950 ng of alkylated RNA using the RiboCop (HMR) kit (as part of the CORALL Total RNA-Seq library Prep Kit, Lexogen 146.24) following the manufacturer's instructions, except for increasing the volume for RNA elution to 16 µl. Removal of rRNAs was confirmed using the RNA 6000 pico kit (Agilent, 5067-1413) on a Bioanalyzer 2100 (Agilent). To prepare cDNA libraries spanning the whole transcript length, the CORALL Total RNA-Seq library Prep Kit (Lexogen 146.24) was used, which includes reverse transcription with Displacement Stop primers. 10 µl of the rRNA depleted

RNA were used as input material and the final library was amplified by 14 PCR cycles. DNA concentration and molarity were determined by Qubit dsDNA HS Assay Kit (Thermo Fisher, Q33230) and Bioanalyzer High Sensitivity DNA Kit (5067-4626) on a Bioanalyzer 2100 (Agilent), respectively. Libraries were pooled and sequenced using the NextSeq 500/550 High Output Kit v2.5 (Illumina, 20024907). The sequencing run was single end with 75 bp reads.

##### **RNA harvest for sequencing from patient-derived xenografts**

Patient-derived xenograft samples were taken from storage at -80°C and broken apart in liquid nitrogen using a mortar and pestle. The tumor fragments were then weighed in as to use at least 20 mg of tumor tissue and homogenized in either TRIzol reagent using a rotor stator instrument or crushing in the mortar and pestle with buffer from the AllPrep DNA/RNA Mini Kit (Qiagen, 80204). Both RNA and DNA were extracted either using the TRIzol (Invitrogen, 15596018) or AllPrep kits. Afterwards 500 ng of RNA per sample was processed using the RiboCop kit (Lexogen, 037) for rRNA removal.

##### **Library preparation and sequencing of ChIP-Rx-Seq and TOP1 CAD-Seq**

DNA from ChIP was quantified with the Qubit dsDNA HS Assay Kit (Thermo Fisher, Q33230). Sequencing libraries were created according to the ThruPLEX DNA-seq kit protocol (Takara, R400676). Size selection was performed in the range of 200-700 bp with AMPure XP beads (Beckman, A63880) and confirmed using the Agilent High Sensitivity DNA Kit (Agilent, 5067-4626) on the Agilent 2100 Bioanalyzer. Libraries were pooled and sequenced using the NextSeq 500/550 High Output Kit v2.5 (Illumina, 20024906). The sequencing run was Single End and Dual Index with 75 bp reads.

##### **Library preparation and sequencing of RNA-Seq**

RNA extracted from xenografts or cultured cells was quantified with the Qubit RNA nano Assay Kit (Thermo Fisher, Q33230). For the Bo99 treated with Irinotecan, JQ1 and Irinotecan+JQ1, library preparation and sequencing was performed by Novogene. For all other sequencing, libraries were created according to the CORALL kit protocol (Lexogen, 095). Library preparation success was confirmed using the Agilent High Sensitivity DNA Kit (Agilent, 5067-4626) on the Agilent 2100 Bioanalyzer. Libraries were pooled and sequenced using the NextSeq 500/550 High Output Kit v2.5 (Illumina, 20024906). The sequencing run was Single End and Dual Index with 75 bp reads for the *in vitro* RNA-Seq from cultured cells, while it was Paired end and Dual Index with 75 bp reads each for the one from xenografts.

##### **ChIP-Rx-Seq data analysis**

For the ChIP-Rx-Seq experiments, the generated fastq files were quality controlled with FastQC and MultiQC (107), trimmed with cutadapt, aligned to the human hg38 and mouse mm10 reference genomes with bowtie2 (108), deduplicated, sorted and indexed using samtools (109) and Picard (110). BigWig files for visualization were generated using Deeptools (111) using -scaleFactor option for spike-in normalization. BigWig files of individually spike-in normalized replicas were merged using UCSC bigwigmerge and bedGraphToBigWig for visual representation. The profiles of short reads average distribution near TSSs and along normalized gene bodies were generated by ngs.plot (112), custom R scripts, Bioconductor (113) packages and ggplot2. Spike-in normalization of average profiles was performed by multiplying the total number of reads by a factor inversely proportional to mouse spike-in unique deduplicated reads and an average of two normalized replicas was used for average profile visualization. Only the protein coding genes from Ensembl 76 (114) database were used for generating the profiles. The profiles were smoothed using a local polynomial regression algorithm. The top 10000 expressed genes were determined based on the RNA-Seq from *in vitro* Bo103 cells treated with DMSO. Box plots were generated in R. Peaks were called from the H3K27ac- and H3K36me-ChIPs using MACS2 (115). Using the ROSE algorithm (42, 102), enhancers were then predicted from the H3K27ac peaks called in the untreated (DMSO) condition.

##### RNA-Seq data analysis

For the modified RNA-Seq and SLAM-seq experiments, the fastq files were prepared the same way as for the ChIP-Rx-Seq data, but the RNA-Seq reads were aligned to hg38 or mm10 reference genome using the splice-aware aligner HISAT2 (116). For deduplication, umi\_tools (117) was used which allows for differentiation between PCR duplicates and naturally occurring deduplication due to the unique molecular identifier (UMI) added through the library preparation by CORALL kit (Lexogen, 095). For the Bo99 data treated with single or combined drugs, deduplication, sorting and indexing were performed using samtools (109) and Picard (110). Reads mapping to both the human and mouse genome were excluded by aligning to one genome first and then using the unmapped reads of that run for alignment with the other genome (human or mouse). Aligned reads were split into two files containing either exonic or non-exonic reads. This was done by first generating a .bed-file of the exon annotations from a .gtf file of Homo sapiens transcriptome assembly release 36 (respectively release 25 for Mus musculus) from GenCode (118) using the grep and gtf2bed from the BEDOPS toolkit (119) commands in the shell. Using this .bed-file, the names of exonic reads were filtered from the respective .bam file using bam2bed and bedops also from BEDOPS, as well as awk in the shell. Finally, using Picard tools' command FilterSamReads, the .bam file was filtered to either include or exclude the list of reads overlapping exons generating two separate .bam-files containing exonic or non-exonic reads. To determine general gene expression level, RNA reads were counted using the featureCounts command from the Subread package (120). Subsequently, the read counts from the untreated (DMSO) condition *in vitro* were used to estimate general gene expression levels by averaging the FPKMs of triplicates. Bigwig files for RNA-Seq visualization were generated using the bamCoverage command from the Deeptools suite (111) after .bam-files had been merged for the *in vitro* RNA-Seq using samtools merge (109).

For differential expression analysis the R package "Index" (121) which is based on the "edgeR" package was used to account for differential expression in both exonic and intronic reads. Exons and introns were counted separately using the filtered .bam files (exonic, non-exonic) instead of assuming the intronic read count based on the difference between reads assigned to the entire gene body and the exons. Significantly differentially expressed genes (adjusted p-value < 0.05 and Log<sub>2</sub> fold change > 1 or < -1) were used for downstream analysis. For visualization of dependence of Log<sub>2</sub> fold change on length, lowly expressed genes (in either exonic or non-exonic reads) were excluded before calculating the moving average of the Log<sub>2</sub> fold-change with a window of 250 genes below and above after ordering the genes by length. To create heatmaps the R package “pheatmap” was used. The filtered non-exonic read files were also used for visualization of read distribution along the gene body and downstream of the TES (non-exonic read distribution or NERD plots) *via* ngs.plot. The ngs.plot output was replotted in R and loess smoothed using the plot-default function. By applying a linear regression along the gene body, the so-called NERD index was calculated which allowed for determination of change of RNA coverage along the gene body and its comparison between treatment conditions. GSEA pre-ranked analysis (GseaPreranked) was performed with software version 4.2.3, using default settings except for “Collapse dataset to gene symbols” set to “No collapse”. Prior to analysis, a list was calculated with each gene assigned a Log<sub>2</sub> p-value and ranked assigning a score based on the false discovery rate (FDR). The ranked genes are annotated according to the gene sets from Gene Ontology (GO) Biological process, selected from the available in “Gene sets database”. Gene sets identified as significant FDR < 0.25 with GSEA.

To detect readthrough and downstream of gene (DoG) transcripts, the algorithm ARTDeco (122) was used on the merged .bam-files. To generate a more reliable set of readthrough genes, the default parameters were changed by extending the minimum length of initially detected DoGs to 12 kb as well as the DoG window size to 2 kb. After DoG detection, a set of high stringency DoG was filtered from the DoGs detected after SN38+JQ1 treatment by counting the RNA-Seq reads covering the detected DoG regions split into windows of 5 kb for 100 kb downstream of the DoG transcript generating gene’s transcription end site (TES) *via* featureCounts. The DoG genes were then filtered for those with a high level of readthrough (CPM > 1.5) 30-45 kb downstream of the TES. From the DoGs detected after SN38+JQ1 treatment *in vitro* a list of non-DoG genes with similar length and expression in this treatment condition was generated. The Venn diagram showing the overlap between *in vivo* and *in vitro* detected DoGs was made using the “Vennerable” package in R.

For analysis of nascent reads via the SLAM-seq method *in vitro*, the established pipeline slamdunk (123) was used. Since the sample preparation deviated from the standard SLAM-seq method, for instance in the library preparation, the pipeline was adapted accordingly. Deduplication via UMI\_tools (117) was performed after alignment via slamdunk map to the hg38 reference genome. After deduplication, slamdunk filter was run with a reference of the entire length of human genes as opposed to just the 3' UTRs, followed by single nucleotide polymorphism (SNP) detection via slamdunk snp. Nascent reads with T > C transitions were then separated from background reads using the alleyoop read-separator command from the slamdunk pipeline. Nascent reads determined via T > C conversions were then visualized using ngs.plot as described above.

##### Repli-seq data analysis

For the Repli-seq data, the generated fastq files were treated the same way as for the ChIP-Rx-Seq files. However, after alignment and deduplication, the established protocol for Repli-seq analysis (59) was followed to determine read coverage in early and late S-phase and the replication timing resulting from the ratio between the two (E, L and E/L respectively). For visualization of replication timing downstream of the TES or at early and late replicated regions of DoGs and non-Dogs, a custom-made python script was developed, before plotting the ensuing data in R with the default plotting function. For determination of early- and late replicated regions the algorithm RepliScan was used (61). The boundaries between those detected regions were then used to determine the distance from the TES of DoG genes detected in SN38+JQ1 and the respective list of non-DoG genes in early regions. Chromosome 3 was excluded from this analysis as there was copy number variation in the 3q arm which interfered with replication timing calling. H3K36me peaks in late replicated regions were determined by taking the MACS2 called peaks and intersecting it with the determined late replicated regions via the intersect command from the BEDTools suite (124).

##### **Graphical design**

Models for inhibitor function, readthrough and replication stress induced by the treatment were created with BioRender.com

##### **Gene Ontology (GO) enrichment analysis**

GO enrichment analysis of biological processes was performed using the software PANTHER Classification System provided by GO web platform. For presentation, representative functional categories were selected based on relevance from the hierarchical clustering tree obtained as an output.

Supplementary Figure 1

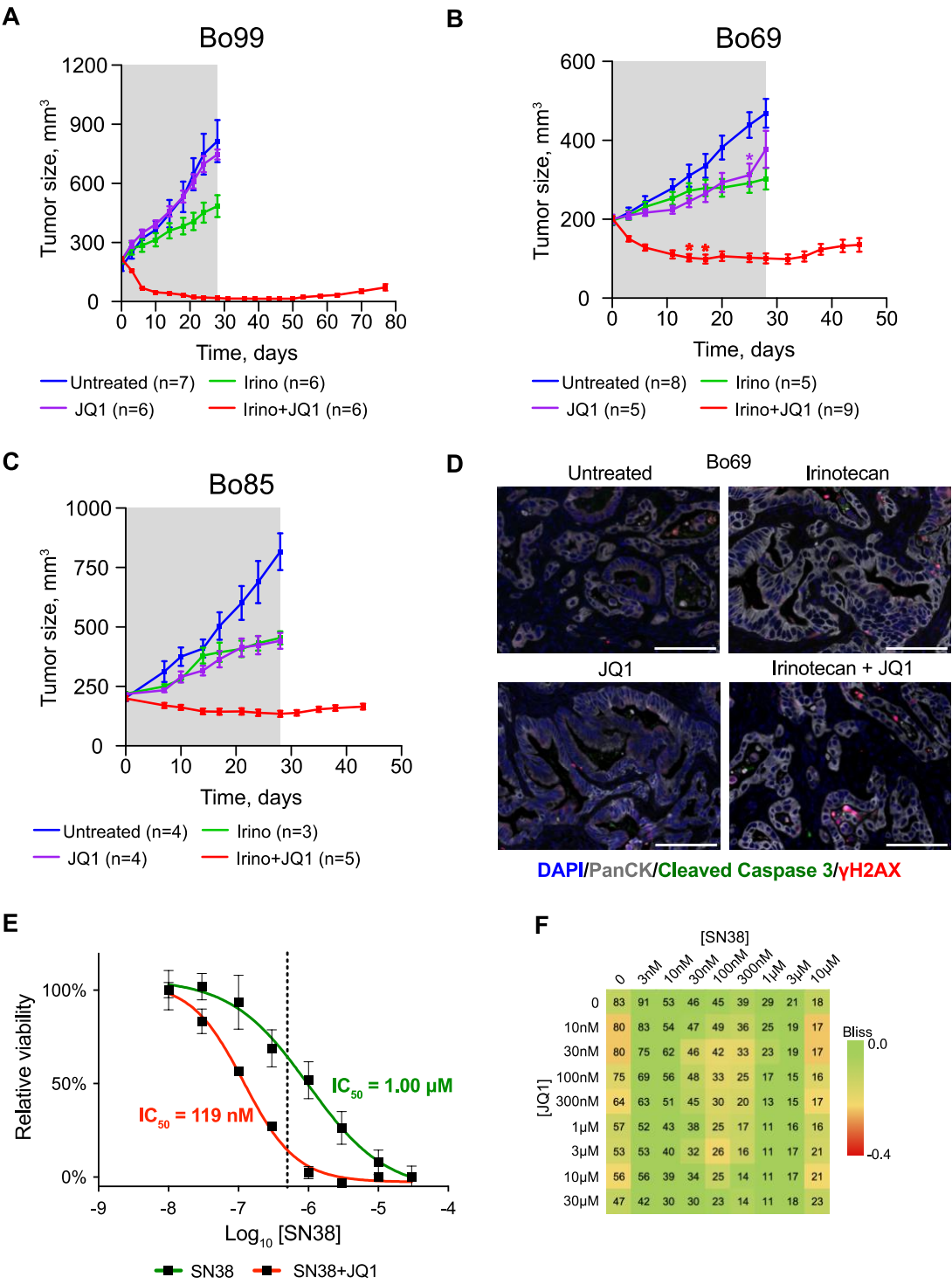

407
408

**Fig. S1. (A-C)** Primary responses observed within the 28 days treatment interval (grey shades area) for the panNEC model (A) Bo99, and the PDAC models (B) Bo69 and (C) Bo85 treated with Irinotecan (15 mg/kg, three times weekly, every second week) and JQ1 (50 mg/kg, daily) by i.p. injection, alone or in combination in comparison to untreated controls (cohort numbers shown). Growth curves are derived from mean values ± SEM (error bars). Each asterisk represents a mouse that was taken out of the treatment cohort (indicated by color) at the indicated time point because of health issues of the animal. **(D)** Representative immunohistochemistry images of Bo69 PDX tumor sections treated for 5 days with Irinotecan and/or JQ1 stained for DNA (DAPI, blue), PanCK (grey), cleaved caspase 3 (green) and γH2AX (red). Scale bar represents 100 μm. **(E)** Dose response curves showing cell viability 48 hr after treatment with 10 nM-30 μM SN38 ± 1 μM JQ1 (n=3). IC<sub>50</sub> values with 95% confidence interval – SN38: 1.00 μM (0.53-1.89 μM); SN38+JQ: 119nM (78-182 nM). **(F)** Checkerboard assay of cultured hTERT-HPNE cells treated with increasing concentrations of SN38 and JQ1 in combination as indicated. The percentage of confluency after treatment is denoted by the numbers in the squares. Synergy was determined using the delta Bliss model of additivity with lower, more negative values showing stronger synergy, (visualized by red/green color coding). Representative checkerboard of n=2.

Supplementary Figure 2

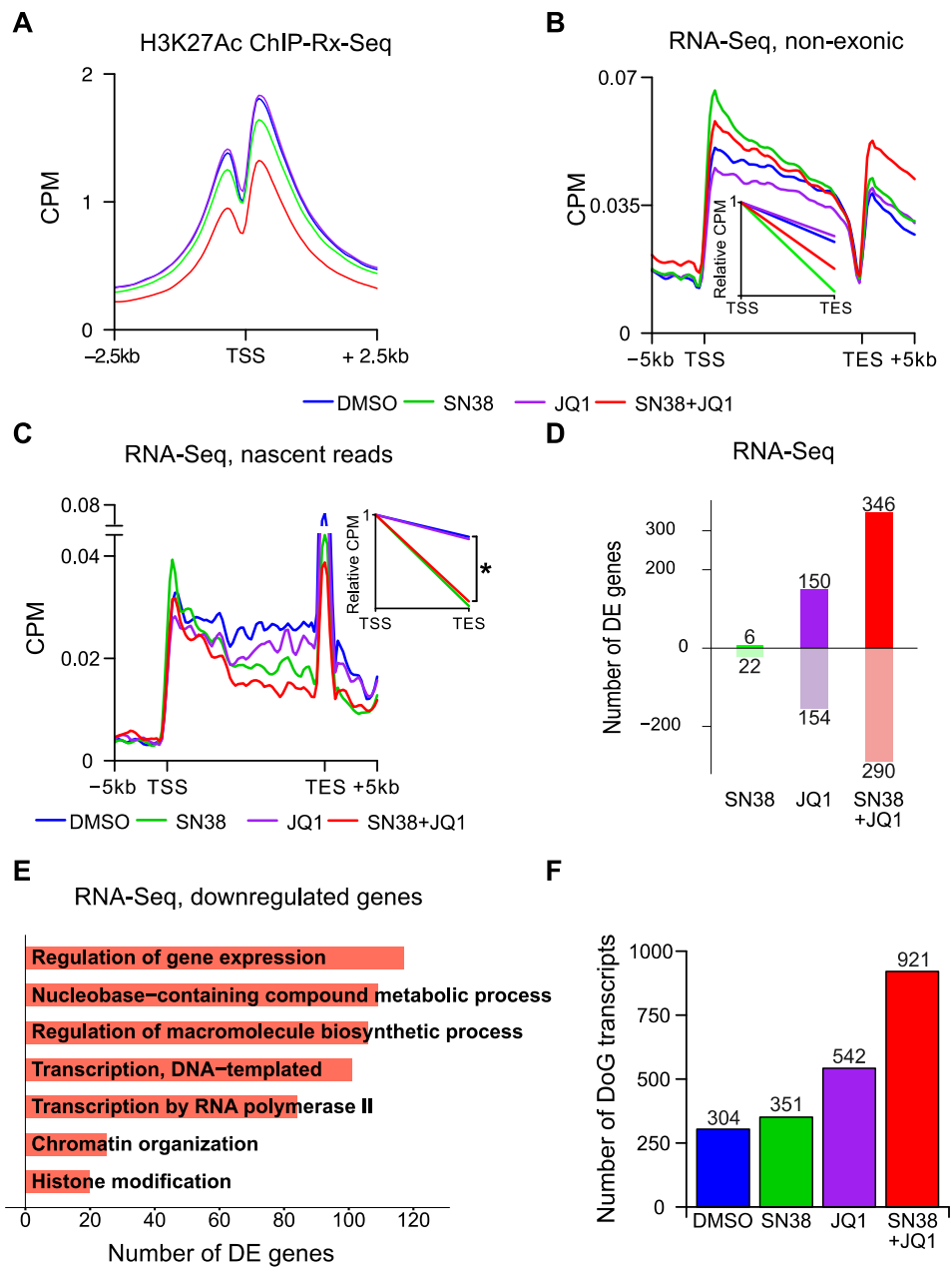

**Fig. S2. (A)** H3K27ac occupancy at the TSS of the 10,000 most expressed genes (+/- 5 kb) in Bo103 cells after 4 hr. Data are spike-in normalized. Average of biological duplicates. **(B)** Non-exonic RNA-Seq reads from Bo103 cells plotted between the TSS and TES of protein-coding genes after 4 hr treatment. Inset shows the gradient of the linear regression in between TSS and TES (NERD index). Average of biological triplicates. **(C)** SLAM-seq nascent (containing T > C transition) reads from Bo103 cells plotted between the TSS and TES of protein-coding genes after 4 hr of treatment. Inset shows the gradient of the linear regression in between the TSS and TES. Spike at TES due to all protein-coding genes having exonic region prior to TES, thus resulting in relative elevated 3' read count. Average of biological triplicates. **(D)** Number of significantly differentially expressed genes based on exonic reads of RNA-Seq (hereafter indicated as RNA-seq) in Bo103 cells treated with SN38, JQ1, or SN38+JQ1 vs. DMSO-treated cells. **(E)** Gene ontology analysis of significantly downregulated genes ($\text{Log}_2\text{FC} < -1$, adjusted $p$-value $< 0.05$) from RNA-Seq of Bo103 cells treated with SN38+JQ1 vs. DMSO-treated cells. **(F)** Number of detected DoG transcript producing genes after treating Bo103 cells with DMSO, SN38, JQ1, or SN38+JQ1.

Supplementary Figure 3

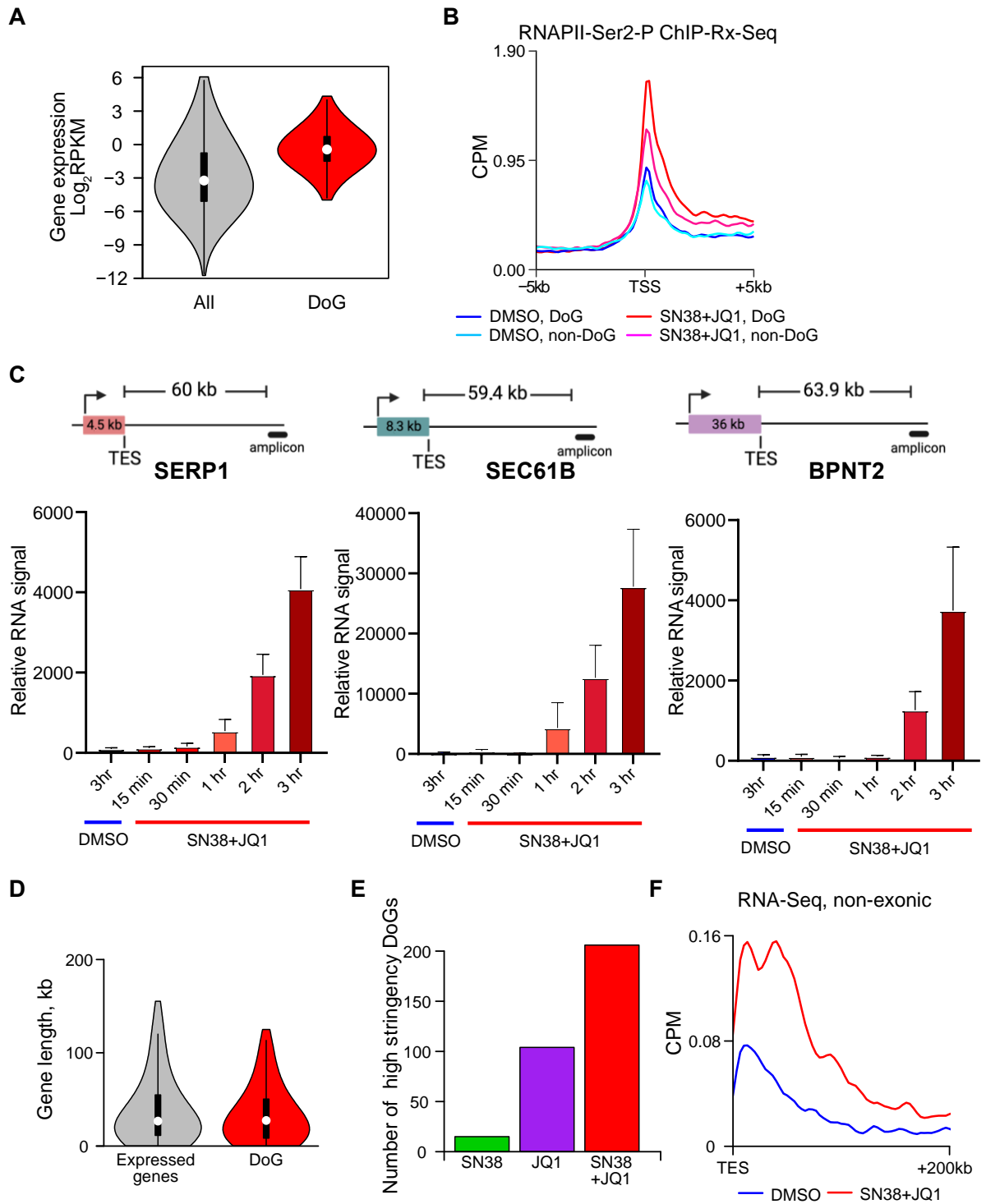

445

446

**Fig. S3.** (A) Violin plot of gene expression level of all genes compared to DoG producing genes; outliers are excluded. (B) RNAPII-Ser2-P occupancy around TSS of DoG and non-DoG genes in Bo103 cells. Data are spike-in normalized. Average of biological duplicates. (C) Top. Schematic showing distance between gene TES and qPCR amplicon. Bottom. Readthrough transcription downstream of DoG genes SERP1, SEC61B and BPNT2 after 15 min up to 180 min of SN38+JQ1 exposure (n=3, relative to DMSO control, error bars represent standard deviation). (D) Violin plot of gene length of all expressed genes compared to DoG producing genes; outliers are excluded. (E) Number of DoG transcripts producing genes of Bo103 cells after filtering for high coverage above untreated levels 30-45 kb downstream of the TES. (F) Non-exonic RNA-Seq reads from Bo103 cells plotted in the region 200 kb downstream of the TES of the high-stringency DoG genes in untreated (DMSO) or treated (SN38+JQ1) conditions. Average of biological triplicates.

#### Supplementary Figure 4

**A**

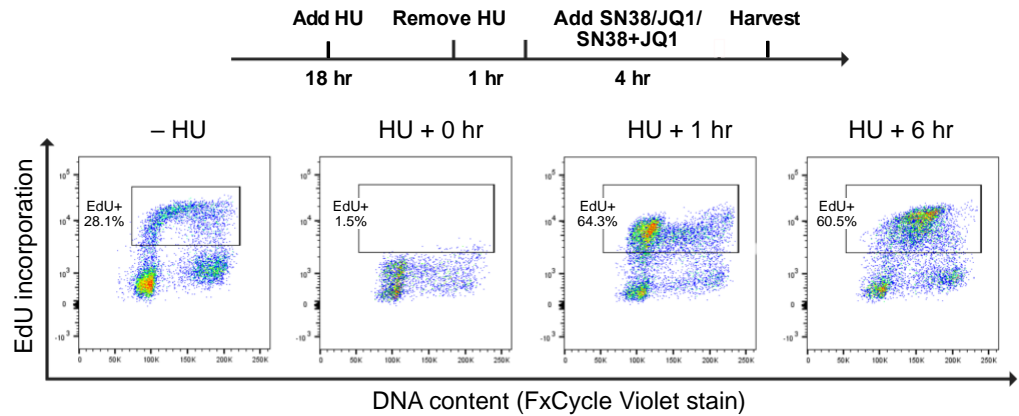

**B**

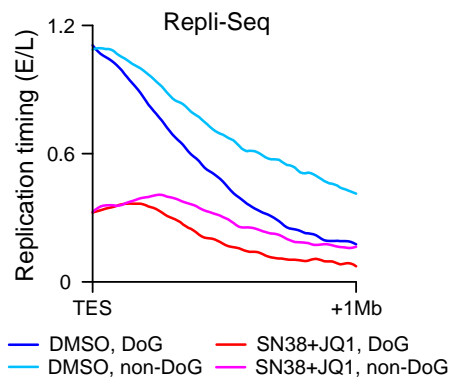

**Fig. S4. (A)** Scheme of S-phase synchronization by Hydroxyurea (HU) for Figure 4A (top) confirmed by flow cytometry (bottom). Representative graphs of multiple timepoints showing the DNA content vs. EdU signal intensity, demonstrating that the majority of cells remain in S-phase up to 6 hr after HU removal. **(B)** Replication Timing determined from Repli-seq in Bo103 cells plotted 1 Mega base pairs (Mb) downstream of the TES of DoGs and non-DoGs in untreated (DMSO) or SN38+JQ1-treated conditions. Greater slope gradient of dark blue (DoG DMSO) vs. light blue (non-DoG DMSO) suggests that regions downstream of DoG genes, on average, transition from early to late replication timing sooner than non-DoG genes. Average of biological duplicates.

**A**

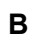

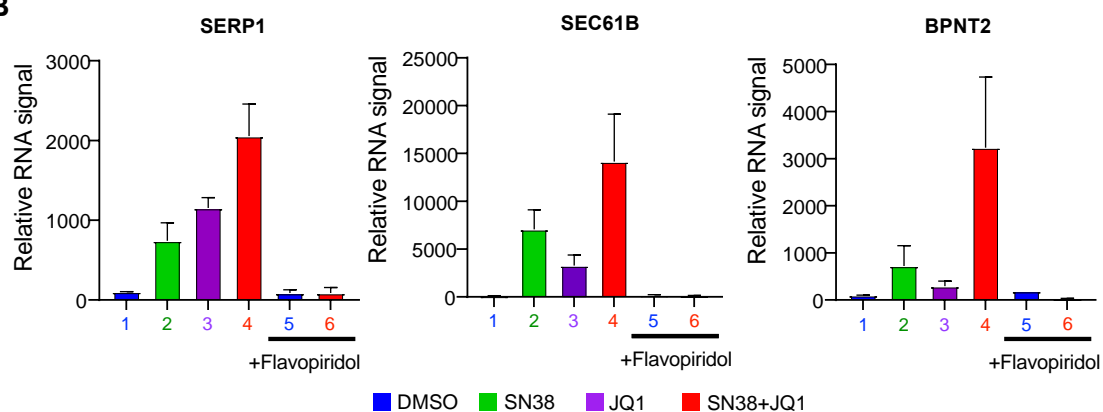

18

Supplementary Figure 6

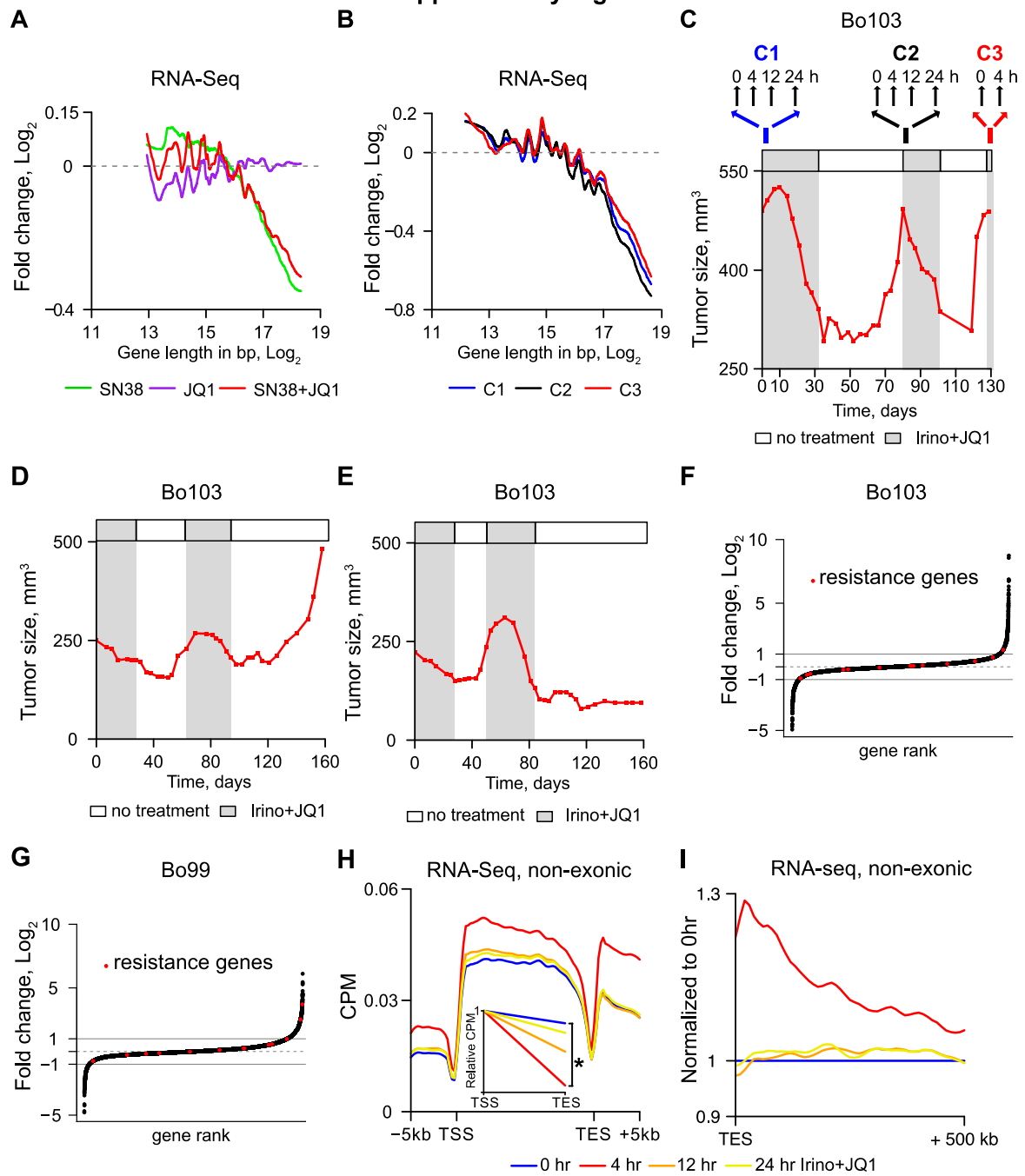

**Fig. S6.** (A) Moving average of fold-change ($\text{Log}_2$) from RNA-Seq reads of treated (SN38, JQ1, or SN38+JQ1) vs. untreated Bo103 cells. Fold-change is plotted against the gene length ($\text{Log}_2$). Average of biological triplicates. (B) Moving average of fold-change ($\text{Log}_2$) of RNA-Seq reads from Bo103 PDX treated with Irinotecan+JQ1 vs. untreated. Fold-change is plotted against the gene length ($\text{Log}_2$). Drug cycles 1, 2 and 3 = C1, C2 and C3. Average of biological triplicates. (C) Top. Dosing and harvesting schedule. Bottom. Representative growth curve of an individual Bo103 PDX tumor, subjected to 3 cycles of treatment with Irinotecan+JQ1. (D-E), Representative growth curves of additional individual Bo103 PDX tumors, subjected to 2 cycles of treatment with Irinotecan+JQ1. (F-G), Ranked list of genes sorted and plotted by $\text{Log}_2$ fold change between C3 0hr and C1 0hr timepoints for (F) Bo103 and (G) Bo99 PDX tumours. Genes associated with resistance to Irinotecan or JQ1 treatment (Supplementary Table S6) highlighted in red. (H) Non-exonic RNA-Seq reads from Bo103 PDX plotted between TSS and TES of protein-coding genes. Inset shows the slope of the curves between TSS and TES. Average of biological triplicates. \*: $p < 0.05$, Student's t-test. (I) Non-exonic RNA-Seq reads of Bo103 PDX plotted in the region 500 kb downstream of the TES of protein-coding genes. Data are expressed as counts per million (CPM) and normalized to the corresponding CPM values at 0 hr.

### Supplementary Figure 7

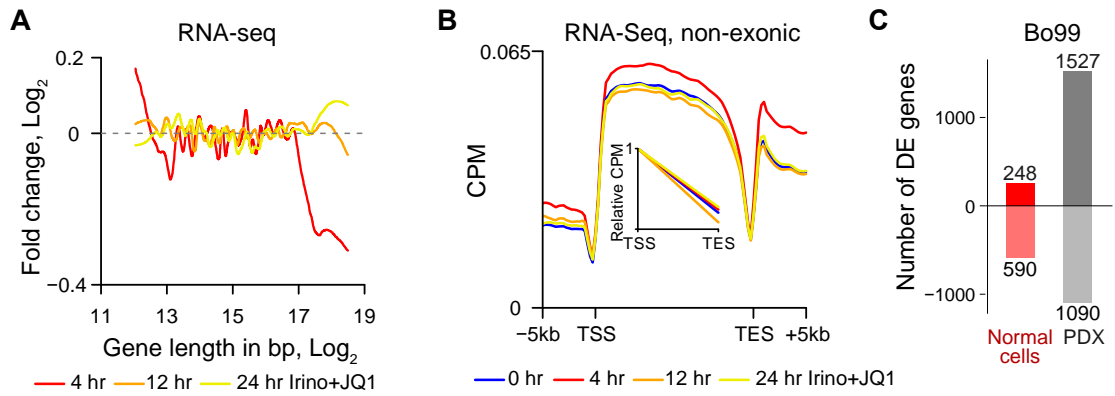

**Fig. S7.** (A) Moving average of fold-change ($\text{Log}_2$) of RNA-Seq reads from normal mouse cells derived from the Bo103 PDX in untreated vs. treated conditions (Irinotecan+JQ1 for 4, 12 and 24 hr). Fold-change is plotted against the gene length ($\text{Log}_2$). Average of 3-4 tumors per condition. (B) Non-exonic RNA-Seq reads from normal mouse cells derived from the Bo103 PDX treated with Irinotecan+JQ1 for different times as indicated. Data are plotted between TSS and TES of protein-coding genes. Inset shows the slope of the curves between TSS and TES (NERD index). Average of 3-4 tumors per condition. No significant difference between NERD indexes is detected. (C) Bar plot representing the number of statistically significant (adjusted $p$-value $< 0.05$) differential expressed up- and down-regulated genes in Bo99 PDXs and normal mouse cells upon treatment with Irinotecan+JQ1 for 4 hr.

**Supplementary Table Legends (provided as Excel files)**

**Table S1.**

PDAC subtype classification, KRAS mutation and MYC expression data for the PDAC and panNEC PDX models.

**Table S2.**

Differential expression analysis from *in vitro* Bo103 RNA-Seq with SN38, JQ1 and SN38+JQ1 treatments.

**Table S3.**

Differential expression analysis from *in vivo* Bo99 RNA-Seq with Irinotecan, JQ1 and Irinotecan+JQ1 treatments.

**Table S4.**

Differential expression analysis of human and mouse exonic reads from *in vivo* Bo99 RNA-Seq with Irinotecan+JQ1 timecourse treatments.

**Table S5.**

Differential expression analysis of human and mouse exonic reads from *in vivo* Bo103 RNA-Seq with Irinotecan+JQ1 timecourse treatments.

**Table S6.**

List of genes associated with resistance to either TOP1 or BRD4 inhibitions, highlighting differential expression ($\Delta\text{Log}_2$ fold-change) in Bo99 and Bo103 between C1 and C3 0 hr timepoints and mean expression levels ($\text{Log}_2\text{FPKM}$). "Regulation" column indicates whether gene is reported to be upregulated or downregulated in resistant cells.
